# Age-dependent accumulation of RAD51 on non-damaged chromosomes prevents chromosome segregation in mammalian oocytes

**DOI:** 10.64898/2026.03.18.712809

**Authors:** Masaru Ito, Shou Soeda, Tomo Kondo, Asako Furukohri, Midori Kajitani, Rina Ogata, Miho Ohsugi, Akira Shinohara

## Abstract

RAD51 is targeted to single-stranded (ss)DNA for homologous recombination and DNA replication fork homeostasis. However, the physiological consequences of RAD51 binding to intact double-stranded (ds)DNA, which is tightly limited *in vivo*, remain elusive. Here we revealed an intrinsic property of RAD51 to bind chromosome axes where cohesin and condensin bind, which is actively suppressed by FIGNL1 AAA+ ATPase. In *Fignl1*-deficient mouse oocytes, an age-dependent RAD51 accumulation with little DNA damage leads to improper chromosomal localization of condensin II and topoisomerase II, failure in chromosome condensation with massive chromosome entanglement, and meiosis I arrest. We propose that promiscuous RAD51 binding to non-damaged chromosomes, which is prevented by a RAD51 remodeler, is a unique type of chromosomal pathology associated with genome instability.

## Introduction

RAD51 recombinase plays a pivotal role in homologous recombination (HR) and protection and restart of stalled replication forks to maintain genome stability (*1*). Although RAD51 forms nucleoprotein filaments on both single-stranded (ss) and double-stranded (ds) DNA with similar affinities *in vitro*, RAD51 is preferentially targeted to ssDNA at the sites of DNA double-strand breaks (DSBs) and collapsed replication forks *in vivo*.

AAA+ ATPase FIGNL1 and its binding partner FIRRM function in dissociating RAD51 from both ssDNA and dsDNA. DNA repair is defective in mouse and human cells deficient for *FIGNL1* and *FIRRM*, and their knockouts (KO) cause lethality of mouse embryonic stem (ES) cells and early embryos (*2–10*). Germ-cell specific conditional knockouts (cKO) of *Fignl1* and *Firrm* in male mice compromised meiotic recombination and homolog synapsis, resulting in the arrest of meiosis progression before pachynema in spermatocytes and infertility (*3, 8, 9*). These indicate the critical role of FIGNL1-FIRRM-mediated RAD51 dissociation from DNA during HR in both somatic and meiotic cells.

A prominent feature of *Fignl1* and *Firrm* cKO spermatocytes is aberrant RAD51 binding to intact meiotic chromatin independently from programmed meiotic DSBs (*3, 8*), which is not associated with DNA damage and repair. However, the impacts of persistent RAD51 binding to intact dsDNA on chromosomal events remained elusive. Here, we show an essential role of FIGNL1-mediated removal of RAD51 from chromosome axes for chromosome condensation and segregation in mouse oocytes.

## Results

### *Fignl1*-deficient oocytes accumulate nuclear RAD51 and are arrested at meiosis I

Sexually dimorphic response to mutations in RAD51 regulators in mammalian meiosis (*11–14*) prompted us to assess the impacts of *Fignl1*-deficiency on oogenesis. To deplete FIGNL1 specifically at the onset of meiosis in mouse oocytes, we used *Fignl1^flox/Δ^*mice harboring a *Stra8-Cre* transgene (*15*). *Fignl1^flox/Δ^ Stra8-Cre^+^,* hereafter referred to as *Fignl1* cKO, adult female mice bred to wild-type male mice produced smaller litters relative to control *Fignl1^+/+^ Stra8-Cre^+^* mice (**Fig. S1A**). Despite a significant reduction of litter size, *Fignl1* cKO females had ovaries that were similar in size and oocyte numbers to controls (*Fignl1^+/+^ Stra8-Cre^+^* and/or *Fignl1^flox/Δ^*) (**Fig. 1A and B, and Fig. S1B**).

**Fig. 1.**
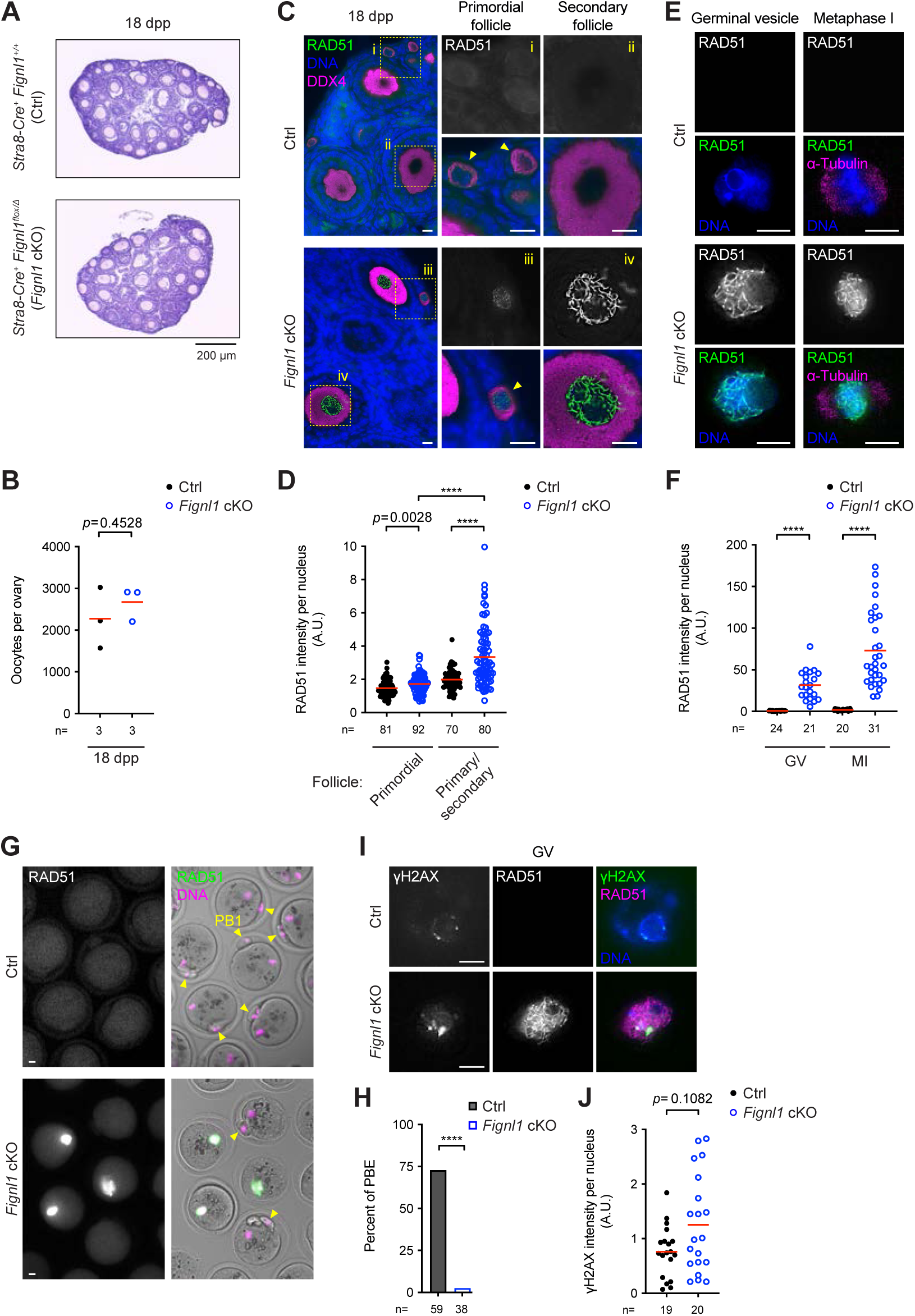
Meiosis-I arrest with aberrant RAD51 accumulation and efficient DNA damage repair in *Fignl1*-deficient oocytes. (A) Representative images of hematoxylin and eosin (HE&E)-stained ovary sections from *Fignl1^+/+^ Stra8-Cre^+^* (Ctrl) and *Fignl1^flox/Δ^ Stra8-Cre^+^*(*Fignl1* cKO) mice at 18 day post-partum (dpp). (B) Oocyte counts of Ctrl and *Fignl1* cKO mice at 18 dpp. The red bars are means. The result of two-tailed unpaired *t*-test is indicated in the graph. Numbers of animals analyzed are indicated below the graph. (C) Representative images of ovary sections stained with DAPI (DNA, blue) and immunostained for RAD51 (white and green) and DDX4 (magenta) from Ctrl and *Fignl1* cKO mice at 18 dpp. Examples of primordial (i and iii) and secondary (ii and iv) follicles are magnified in the right panels. (D) Signal intensities of nuclear RAD51 in Ctrl (black circles) and *Fignl1* cKO (blue open circles) oocytes in ovary sections from 18-dpp animals. The red bars are means. The results of two-tailed Mann-Whitney *U*-tests are indicated in the graph: \*\*\*\**p* ≤ 0.0001. Total numbers of oocytes analyzed are indicated below the graph. (E) Representative images of fixed oocytes stained with Hoechst 33342 (DNA, blue) and immunostained for RAD51 (white in the top panels and green in the bottom panels) and α-tubulin (magenta) at Germinal-vesicle (GV) and metaphase-I stages from Ctrl and *Fignl1* cKO mice. Images are optical slices to show the middle of nuclei. (F) Signal intensities of nuclear RAD51 in fixed Ctrl (black circles) and *Fignl1* cKO (blue open circles) oocytes at GV and metaphase-I stages. The red bars are means. The results of two-tailed Mann-Whitney *U*-tests are indicated in the graph. Total numbers of oocytes analyzed are indicated below the graph. (G) Representative images of fixed oocytes stained with Hoechst 33342 (DNA, magenta) and immunostained for RAD51 (white in the left panels and green in the right panels) at 16 h of culture from Ctrl and *Fignl1* cKO mice. Yellow arrow heads indicate polar bodies (PB1). Images are optical slices to show PBs. (H) Efficiency of polar body extrusion (PBE), an indicator of meiosis I division, in Ctrl (a gray bar) and *Fignl1* cKO (a blue open bar) oocytes after 16 h of culture. The results of Fisher’s exact tests are indicated in the graph: \*\*\*\**p* ≤ 0.0001. Total numbers of oocytes analyzed are indicated below the graph. (I) Representative images of fixed GV oocytes stained with Hoechst 33342 (DNA, blue) and immunostained for γH2AX (white in the left panels and green in the right panels) and RAD51 (white in the middle panels and magenta in the right panels) from Ctrl and *Fignl1* cKO mice. Images are optical slices to show the middle of nuclei. (J) Signal intensities of nuclear γH2AX in fixed Ctrl (black circles) and *Fignl1* cKO (blue open circles) oocytes at the GV stage. The red bars are means. Total numbers of oocytes analyzed are indicated below the graph. Genotypes of indicated animals are: Ctrl, *Fignl1^+/+^ Stra8-Cre^+^*in (A)-(D) and *Fignl1^flox/+^ Stra8-Cre^+^* in (E)-(J); *Fignl1* cKO, *Fignl1^flox/Δ^ Stra8-Cre^+^*. Scale bars in (C), (E), (G), and (I), 10 μm.

Defects in repair of meiotic DSBs and homolog synapsis cause culling of late prophase I oocytes (*16, 17*). Normal postnatal follicle reserves in *Fignl1* cKO female mice suggest that *Fignl1* cKO oocytes complete meiotic prophase I with efficient meiotic DSB repair and homolog synapsis. Surface-spread chromosomes of fetal oocytes at various stages revealed elevated numbers of RAD51 foci throughout meiotic prophase I in *Fignl1* cKO oocytes (**Fig. S1C and D**). Critically, *Fignl1* cKO pachytene oocytes with fully synapsed chromosomes at embryonic day 18.5 (E18.5) retained ∼250 RAD51 foci and some of *Fignl1* cKO dictyate oocytes at 1 day post-partum (dpp) showed ∼800 RAD51 foci. This is in sharp contrast to control *Fignl1^flox/+^ Stra8-Cre^+^* oocytes that showed a reduction of RAD51 focus numbers as prophase I progressed to ∼40 and ∼5 foci in pachytene and dictyate stages, respectively (**Fig. S1C and D**). We note that *Fignl1* cKO ovaries contained two populations of oocytes: one showed aberrant RAD51 accumulation and the other showed nuclear RAD51 levels comparable to control oocytes (**Fig. S1D-F**). The latter is likely derived from oocytes that failed to excise the *Fignl1^flox^* allele, thus *Fignl1^flox/Δ^* cells, which are proficient for *Fignl1* and behaved like control oocytes (**Fig. S1G and H**). Hereafter, we focused on oocytes with high levels of nuclear RAD51 (either focus number or signal intensity) in *Fignl1* cKO mice as *Fignl1*-deficient oocytes otherwise mentioned.

Despite aberrant RAD51 accumulation, repair of meiotic DSBs was efficient. RPA2 is a component of RPA complex that specifically binds ssDNA and thus is a reliable marker of DSB repair (*18*). Although RPA2 focus numbers were higher in *Fignl1* cKO than in control oocytes throughout prophase I, those in most of *Fignl1* cKO oocytes at diplotene and dictyate stages were comparable to control oocytes (**Fig. S2A and B**). The number and distribution of MLH1 foci, which mark crossover sites (*19, 20*), were indistinguishable between *Fignl1* cKO and control pachytene oocytes at E18.5 (**Fig. S2C-E**).

Ovary sections from *Fignl1* cKO mice at 18 dpp showed the presence of primordial, primary, and secondary follicles with higher immunostaining signals of RAD51 in the oocyte nuclei (**Fig. 1C)**, in contrast to nearly background levels of RAD51 in those of control mice. Notably, immunostaining pattern of RAD51 changed from punctate foci in primordial follicles to lines or bundles in primary and secondary follicles, which was accompanied by an increase in nuclear RAD51 signals (**Fig. 1C and D**), suggesting further accumulation of RAD51 in growing follicles in *Fignl1* cKO mice. In contrast, a meiosis-specific RAD51 homolog DMC1, which was also accumulated in *Fignl1* cKO oocytes during early prophase I, showed a reduction of its focus numbers as prophase I progressed to nearly control levels in the dictyate stage at 1 dpp (**Fig. S3A and B**), and none of *Fignl1* cKO oocytes was positive for DMC1 immunostaining at 18 dpp (**Fig. S3C and D**).

Meiotic arrest of oocytes in the dictyate stage and successful growth of preovulatory follicles in *Fignl1* cKO mice led us to analyze later steps of oocyte meiosis. *In vitro* maturation experiments revealed that *Fignl1* cKO oocytes, which showed multiple RAD51 lines with ∼50-fold higher immunostaining signals of nuclear RAD51 than control oocytes at the germinal vesicle (GV) stage, underwent GV breakdown (GVBD) efficiently, and formed metaphase I-spindles (**Fig. 1E and F, and Fig. S1G**). However, 97.4% (37/38) of *Fignl1* cKO oocytes failed to extrude a polar body (PB) (**Fig. 1G and H, and Fig. S1H**). These results indicate that *Fignl1* cKO oocytes complete meiotic prophase I, enter the dictyate arrest and resume meiosis efficiently, but are arrested at meiosis I with persistent accumulation of RAD51 on meiotic chromosomes. The presence of a considerable number of RAD51-negative, *Fignl1*-proficient oocytes in *Fignl1* cKO ovaries (65.6%, 565/861 oocytes from two *Fignl1* cKO mice at 1-2 dpp) could explain a small reduction in fertility of *Fignl1* cKO female mice (**Fig. S1A**).

### *Fignl1*-deficient oocytes show little increase in DNA damage during oocyte growth

Defective DNA repair during the dictyate arrest can also cause oocyte loss (*21*). Large postnatal follicle reserves in *Fignl1* cKO female mice suggest that postnatal *Fignl1* cKO oocytes are proficient for DNA repair. This inference was tested by monitoring phosphorylated histone H2AX (γH2AX) as a marker of DNA damage. In ovary sections from 18-dpp animals, while nuclear γH2AX levels in primordial follicles were 1.2-fold higher in *Fignl1* cKO oocytes than in control oocytes, those in primary/secondary follicles were indistinguishable between *Fignl1* cKO and control oocytes (**Fig. S4A and B**). Consistently, γH2AX levels in *Fignl1* cKO and control oocytes were comparable at the GV stage (**Fig. 1I and J**). These results suggest that residual DSBs that remained unrepaired during prophase I could be repaired during the dictyate arrest, and RAD51 accumulation during the oocyte growth does not accompany increased DNA damage in *Fignl1* cKO oocytes.

### Chromosomes fail to condense and segregate at meiosis I in *Fignl1*-deficient oocytes

Efficient DSB repair and crossover formation during prophase I and the dictyate arrest along with the oocyte growth in *Fignl1* cKO oocytes led us to image live oocytes to uncover the cause of meiotic arrest. Control oocytes showed rapid chromosome condensation and formed 20 bivalents soon after GVBD, which resulted in a reduction of chromosome volume by half of that before GVBD, and successful segregation of homologous chromosomes to extrude the first PBs by 16 h of culture (**Fig. 2A-C and movie S1**). In sharp contrast, in oocytes from *Fignl1* cKO mice, chromosomes did not condense or individualize, with a reduction in chromosome volume by only ∼20% after GVBD, and failed to segregate even after >16 h of culture (**Fig. 2A-C and movie S2**). In those oocytes, some chromosome fragments were occasionally separated from but rapidly incorporated into a chromosome mass (**Fig. 2A and B, and movie S2**). Chromosome spreads of metaphase-I oocytes at 7 h of culture confirmed RAD51 accumulation in oocytes that failed chromosome condensation (**Fig. 2D and E**). An end of each chromosome fragment separated from a chromosome mass in *Fignl1* cKO oocytes was often labelled with anti-centromere antibody (ACA) (**Fig. 2D**). This suggests that spindles pull and can temporally dissociate centromeres from an uncondensed chromosome mass, but failed to segregate homologous chromosomes at meiosis I in *Fignl1* cKO oocytes.

**Fig. 2.**
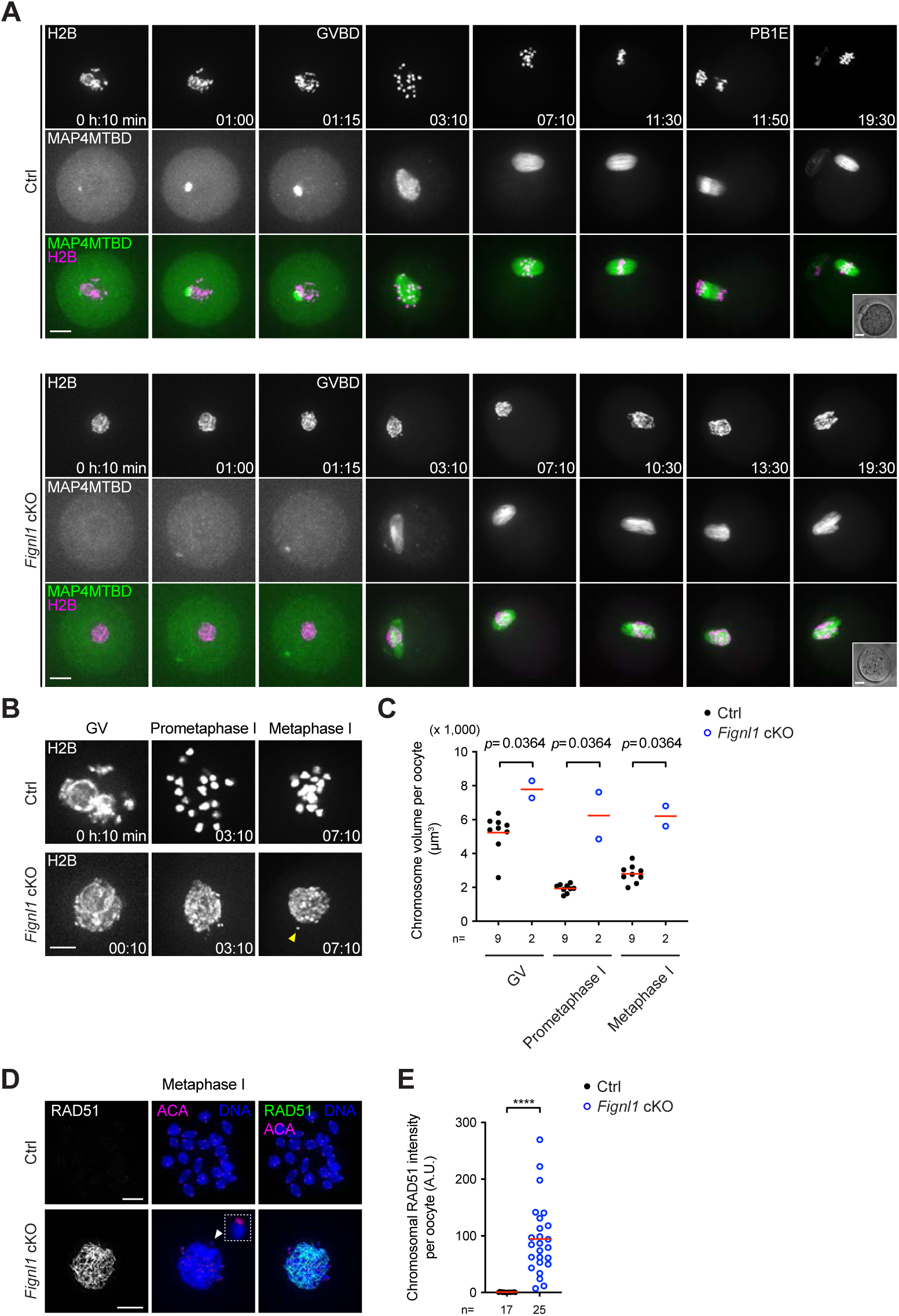
Defective chromosome condensation and segregation at meiosis I in *Fignl1*-deficient oocytes. (A) Representative still images of oocytes microinjected with mRNAs encoding histone H2B–mRFP (white in the top panels and magenta in the bottom panels) and EGFP–MAP4MTBD (white in the middle panels and green in the bottom panels) from Ctrl and *Fignl1* cKO mice. Images are maximal intensity *z*-projections. Time 0 corresponds to the point of dbcAMP removal for meiotic resumption. Insets show bright-field images. PB1E, polar body extrusion. (B) Enlarged view of H2B images in (A) at GV, prometaphase-I and metaphase-I stages. An yellow arrowhead indicates a chromosome fragment separated from a chromosome mass. (C) Chromosome volume of GV, prometaphase-I and metaphase-I oocytes from Ctrl (black circles) and *Fignl1* cKO (blue open circles) mice. The results of two-tailed Mann-Whitney *U*-tests are indicated in the graph. Total numbers of oocytes analyzed are indicated below the graphs. (D) Representative images of chromosome spreads of metaphase-I oocytes stained with DAPI (DNA, blue) and immunostained for RAD51 (white in the left panels and green in the right panels) and centromeres (ACA, magenta) from Ctrl and *Fignl1* cKO mice. A white arrowhead indicates a chromosome fragment containing centromeres separated from a chromosome mass, which is magnified in an inset. (E) Signal intensities of chromosomal RAD51 on chromosome spreads of metaphase-I Ctrl (black circles) and *Fignl1* cKO (blue open circles) oocytes. The red bars are means. The results of two-tailed Mann-Whitney *U*-tests are indicated in the graph: \*\*\*\**p* ≤ 0.0001. Total numbers of oocytes analyzed are indicated below the graphs. Genotypes of indicated animals are: Ctrl, *Fignl1^flox/+^ Stra8-Cre^+^*; *Fignl1* cKO, *Fignl1^flox/Δ^ Stra8-Cre^+^*. Scale bars, 20 μm in (A) and 10 μm in (B) and (D).

### RAD51 accumulation leads to improper chromosomal localization of condensin II and topoisomerase II with compromised topoisomerase II function

To gain mechanistic insights into how *Fignl1*-deficiency causes defective chromosome condensation and segregation, we analyzed chromosomal localization of the condensin complex. The condensin complexes are essential for chromosome condensation in both mitosis and meiosis, and form two distinct types of complexes, condensin I and II, in mammals (*22*). NCAPD3, a component of condensin II that likely plays a primary role at meiosis I (*23, 24*), localized to the entire chromosomes of *Fignl1* cKO metaphase-I oocytes as punctate foci, in sharp contrast to its preferential enrichment on chromatid axes of bivalent chromosomes in control oocytes (**Fig. 3A**).

**Fig. 3.**
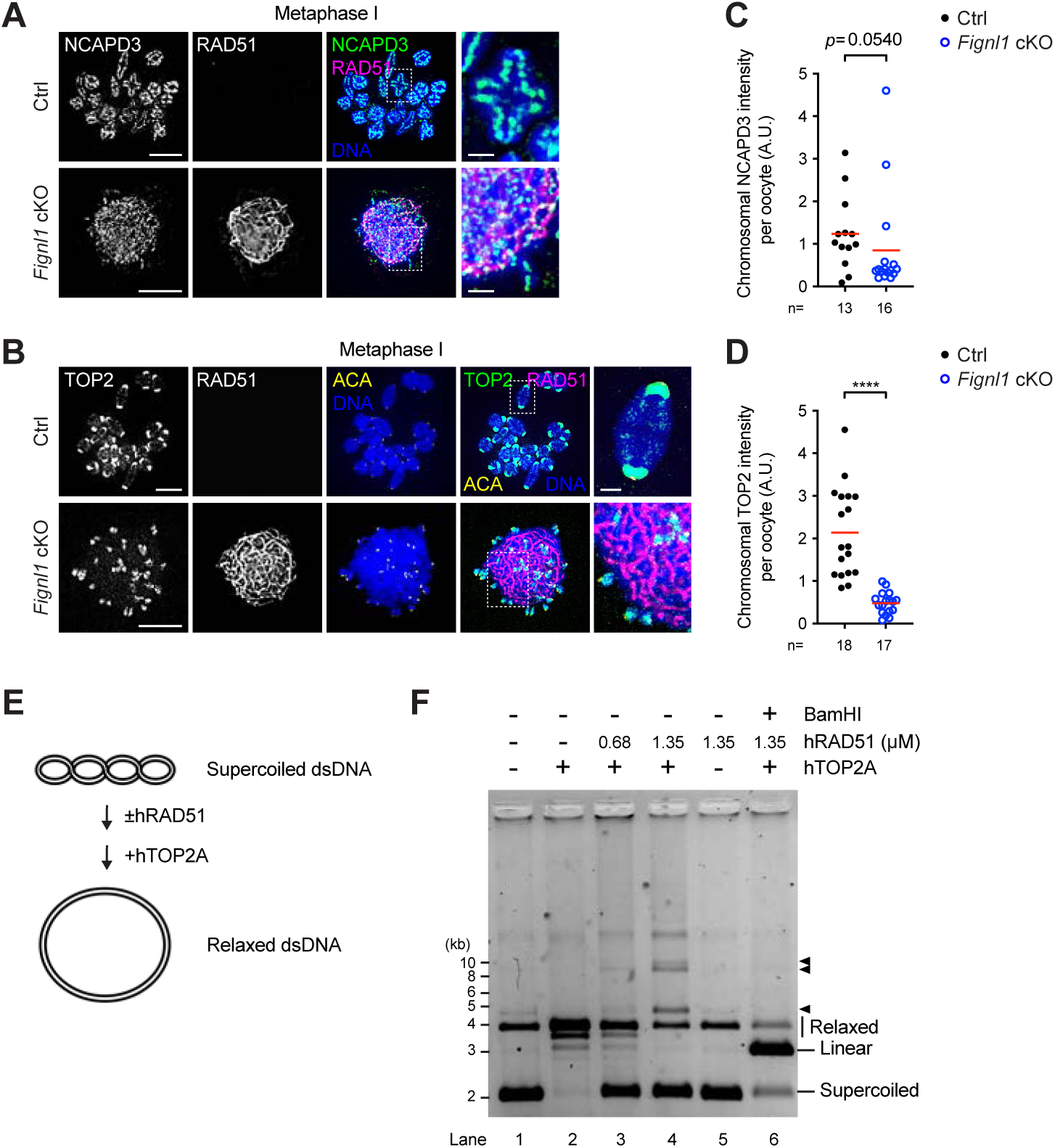
Improper chromosomal localization and function of condensin II and topoisomerase II on RAD51-bound DNA. (A) Representative images of chromosome spreads of metaphase-I oocytes stained with DAPI (DNA, blue) and immunostained for NCAPD3 (white in the left panels and green in the right panels) and RAD51 (white in the left panels and magenta in the right panels) from Ctrl and *Fignl1* cKO mice. (B) Representative images of chromosome spreads of metaphase-I oocytes stained with DAPI (DNA, blue) and immunostained for TOP2 (white in the left panels and green in the right panels), RAD51 (white in the left panels and magenta in the right panels), and centromeres (ACA, yellow) from Ctrl and *Fignl1* cKO mice. (C and D) Signal intensities of chromosomal NCAPD3 (C) and TOP2 (D) on chromosome spreads of metaphase-I Ctrl (black circles) and *Fignl1* cKO (blue open circles) oocytes. The red bars are means. The results of two-tailed Mann-Whitney *U*-tests are indicated in the graphs: \*\*\*\**p* ≤ 0.0001. Total numbers of oocytes analyzed are indicated below the graphs. (E) Reaction scheme of DNA relaxation assay. (F) A representative gel image of DNA relaxation assay showing *in vitro* activity of TOP2A to remove helical tension in supercoiled dsDNA. 3.8 μM in bp of pBluescript II SK+ plasmid was preincubated with and without human RAD51 at indicated concentrations at 37°C for 5 min, and 5 units of human TOP2A was added and further incubated at 37°C for 20 min. BamHI was finally added in the indicated reaction and further incubated at 37°C for 5 min. Migration positions of supercoiled, linear, and relaxed or open circular DNA are indicated. Black arrowheads indicate slower migration products specifically detected or enriched in the presence of RAD51 in the TOP2A reaction (lanes 3 and 4). Genotypes of indicated animals are: Ctrl, *Fignl1^flox/+^ Stra8-Cre^+^*; *Fignl1* cKO, *Fignl1^flox/Δ^ Stra8-Cre^+^*. Scale bars in (A) and (B), 10 μm for full nuclei and 2 μm for magnified panels.

We also analyzed chromosomal localization of topoisomerase II (TOP2), another critical factor for chromosome condensation in both mitosis and meiosis (*24–27*). TOP2 formed punctate foci on chromatid axes with its strong enrichment at centromeric heterochromatin marked by histone H3 lysine 9 trimethylation (H3K9me3) on metaphase-I chromosomes of control oocytes (**Fig. 3B and Fig. S5A**). In *Fignl1* cKO oocytes, although its strong enrichment at centromeric heterochromatin was retained, its punctate foci were more diffusely localized to the entire chromosomes, like NCAPD3 (**Fig. 3B and Fig. S5A**). While chromosomal NCAPD3 levels were comparable between *Fignl1* cKO and control oocytes (**Fig. 3C**), chromosomal TOP2 levels in *Fignl1* cKO oocytes were reduced by half of those in control oocytes (**Fig. 3D**).

TOP2 contributes to chromosome condensation via removal of DNA catenates and knots (*25*). *In vitro* DNA relaxation assay revealed that RAD51 binding to DNA reduced the efficiency of human TOP2A to relax dsDNA supercoils (**Fig. 3E and F, and Fig. S5B**). Qualitatively, DNA molecules with slower migration on the gels specifically detected in the presence of RAD51 (**Fig. 3F and Fig. S5B, arrowheads**) implicates unregulated TOP2 function on RAD51-bound DNA. These results suggest that RAD51 accumulation impedes proper chromosomal localization of the two major factors for chromosome condensation and compromises TOP2 function.

### *Fignl1* deficiency leads to preferential RAD51 accumulation to chromosome axes with cohesin REC8 and condensin II

To examine the spatial relationship between RAD51 accumulation and localization of condensin II and TOP2 in *Fignl1* cKO oocytes, we performed Chromatin immunoprecipitation-sequencing (ChIP-seq) for RAD51 in *Fignl1* cKO mouse spermatocytes (*3*) and compared with published ChIP-seq data for NCAPH2, a component of condensin II complex, in mouse ES cells (*28*). Consistent with specific localization of immunostaining RAD51 foci on chromosome axes in both *Fignl1* cKO spermatocytes (*3, 8*) and oocytes (**Fig. S1C**) during prophase I, RAD51 was clearly enriched at chromosome axes, especially where cohesin REC8 was enriched (**Fig. 4A and Fig. S6A-C**). Notably, NCAPH2 was specifically enriched at where REC8 strongly binds, and higher NCAPH2 enrichment corresponded to higher enrichment of REC8 and RAD51 (**Fig. 4A**). These data indicate that RAD51 preferentially binds chromosome axes, especially where REC8 and condensin II bind, in mouse spermatocytes. We note that RAD51 binding to chromosome axes, which was detectable even in wild-type spermatocytes (**Fig. S6A-C**), was not detected in our previous ChIP-single-stranded DNA sequence (SSDS) (*3*), which specifically detected its binding to ssDNA around meiotic DSB hotspots (**Fig. S6B and C**). This suggests that conventional ChIP for RAD51 preferentially captures its binding to dsDNA, and thus RAD51 binds intact dsDNA at chromosome axis sites.

**Fig. 4.**
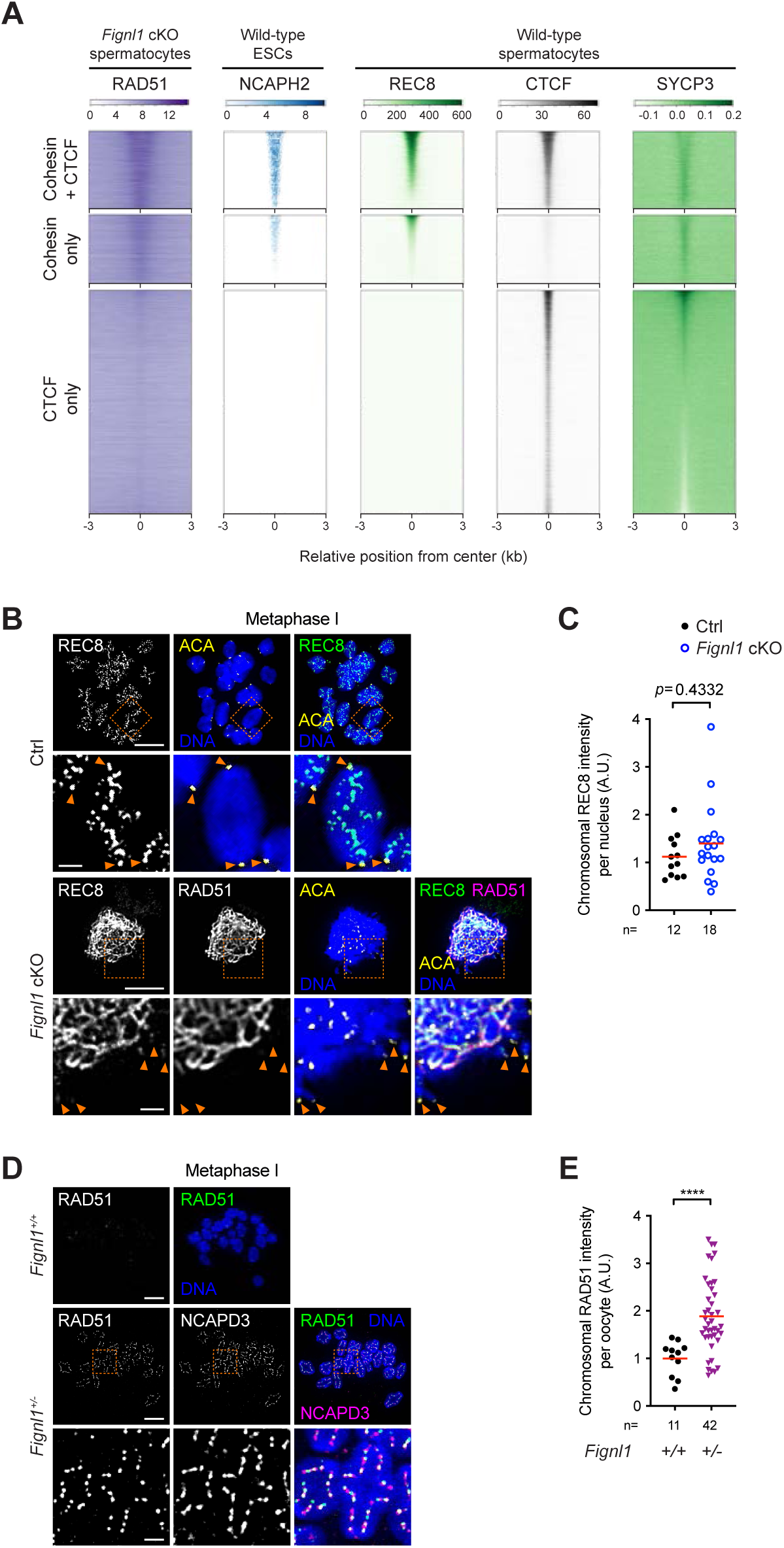
Preferential accumulation of RAD51 to chromosome axis sites where condensin II and cohesin REC8 bind by *Fignl1* depletion. (A) Heatmaps of ChIP signals of RAD51 in *Fignl1* cKO mouse spermatocytes (purple), NCAPH2 in mouse ES cells (blue), and REC8 (green), CTCF (black), and SYCP3 (green) in wild-type mouse spermatocytes around chromosome axes. Chromosome axis sites are classified into three based on enrichment of REC8 and CTCF as our previous study. 7,132 CTCF + cohesin sites, 6,379 cohesin-only sites, and 19,939 CTCF-only sites are ordered by REC8, REC8, and SYCP3 enrichment, respectively, ChIP data for NCAPH2, REC8, CTCF and SYCP3 are from previous studies (*28, 35, 36*). (B) Representative images of chromosome spreads of metaphase-I oocytes stained with DAPI (DNA, blue) and immunostained for REC8 (white in the left panels and green in the right panels), RAD51 (white in the left panels and magenta in the right panels), and centromeres (ACA, yellow) from Ctrl and *Fignl1* cKO mice. Orange arrowheads indicate centromeres. (C) Signal intensities of chromosomal REC8 on chromosome spreads of metaphase-I Ctrl (black circles) and *Fignl1* cKO (blue open circles) oocytes. The red bars are means. (D) Representative images of chromosome spreads of metaphase-I oocytes stained with DAPI (DNA, blue) and immunostained for RAD51 (white in the left panels and green in the right panels) and NCAPD3 (white in the middle panels and magenta in the right panels) from wild-type (*Fignl1^+/+^*) and *Fignl1* heterozygous (*Fignl1^+/-^*) mice. (E) Signal intensities of chromosomal RAD51 on chromosome spreads of metaphase-I *Fignl1^+/+^* (black circles) and *Fignl1^+/-^* (purple triangles) oocytes. The red bars are means. The results of two-tailed Mann-Whitney *U*-tests are indicated in the graphs: \*\*\*\**p* ≤ 0.0001. Total numbers of oocytes analyzed are indicated below the graphs. Genotypes of indicated animals are: Ctrl, *Fignl1^flox/+^ Stra8-Cre^+^*; *Fignl1* cKO, *Fignl1^flox/Δ^ Stra8-Cre^+^*. Scale bars in (B) and (D), 10 μm for full nuclei and 2 μm for magnified panels.

To address whether RAD51 accumulates at REC8-binding sites in *Fignl1* cKO oocytes, chromosomal localization of REC8 was analyzed on chromosome spreads of metaphase-I oocytes. REC8 formed punctate foci primarily localized between sister chromatids of bivalent chromosomes and centromeres in control oocytes (**Fig. 4B**). In contrast, REC8 showed linear immunostaining pattern in *Fignl1* cKO oocytes along with foci at centromeres, with its chromosomal levels comparable to control oocytes (**Fig. 4B and C**). Co-staining with RAD51 revealed substantial colocalization of REC8 and RAD51 lines, supporting the idea that RAD51 preferentially binds REC8-bound chromosome axes in *Fignl1* cKO metaphase-I oocytes.

We note that *Fignl1^+/-^* heterozygous metaphase-I oocytes, which formed normal bivalent chromosome structures, showed higher chromosomal RAD51 levels than wild-type oocytes, and RAD51 formed immunostaining foci primarily on chromatid axes where NCAPD3 localizes (**Fig. 4D and E**). This indicates that FIGNL1 regulates RAD51 dissociation from DNA in a dosage-dependent manner in mouse metaphase-I oocytes and further supports the idea that RAD51 preferentially localizes to where condensin II binds.

### *Fignl1* depletion in growing oocytes causes an age-dependent increase in chromosomal RAD51 and the severity of chromosome condensation defects

Our results suggest that RAD51 accumulation at chromosome axes is a leading cause of defective chromosome condensation and segregation in *Fignl1* cKO oocytes. Coincidence of cohesin REC8 and condensin II binding sites and linear immunostaining pattern of REC8 in *Fignl1* cKO metaphase-I oocytes, which was not seen in control oocytes (**Fig. 4B**), raise the possibility that persistent or altered REC8 localization may impede proper localization of condensin II and TOP2 to meiotic chromosomes. However, this is unlikely because REC8 depletion by TRIM-Away (*29, 30*), which efficiently produced univalent chromosomes in control oocytes (**Fig. S7A**) and decreased chromosomal REC8 levels in *Fignl1* cKO oocytes by ∼40% (**Fig. S7B and C**), had little effects on localization of both NCAPD3 and TOP2 to the entire chromosome mass in *Fignl1* cKO metaphase-I oocytes (**Fig. S7A and D**).

Inability to efficiently deplete RAD51 by TRIM-Away (**Fig. S8A and B**) or remove RAD51 from chromosomes by B02 RAD51 inhibitor (**Fig. S8C and D**) in *Fignl1* cKO oocytes in our hands precluded us from directly assessing the role of RAD51 accumulation in defective chromosome condensation and segregation. Given the increase in nuclear RAD51 levels during the oocyte growth (**Fig. 1C and D**), we set up a time course experiment by depleting *Fignl1* in growing oocytes to address whether chromosomal RAD51 levels correlate with the severity of chromosome condensation defects (**Fig. 5A**). Cre recombinase fused with mouse estrogen receptors (MER Cre MER; MCM) under the promoter of *Dppa3* gene (*Dppa3-MCM*) allows tamoxifen-induced deletion of flox alleles in growing oocytes (*31, 32*). *Dppa3-MCM*-mediated *Fignl1* cKO, hereafter referred to as *Fignl1 D*-cKO, showed a time-dependent RAD51 accumulation in GV oocytes: the frequencies of oocytes with linear RAD51 immunostaining signals were increased from 0% (0/14) at 4 days post-tamoxifen treatment (4 dpt) to 23.8% (5/21) and 81.5% (22/27) at 9 and 18 dpt, respectively (**Fig. 5B and C**), and nuclear RAD51 levels were correspondingly increased in proportion to days post-tamoxifen treatment with little increase in γH2AX levels (**Fig. 5D and Fig. S9A and B**).

**Fig. 5.**
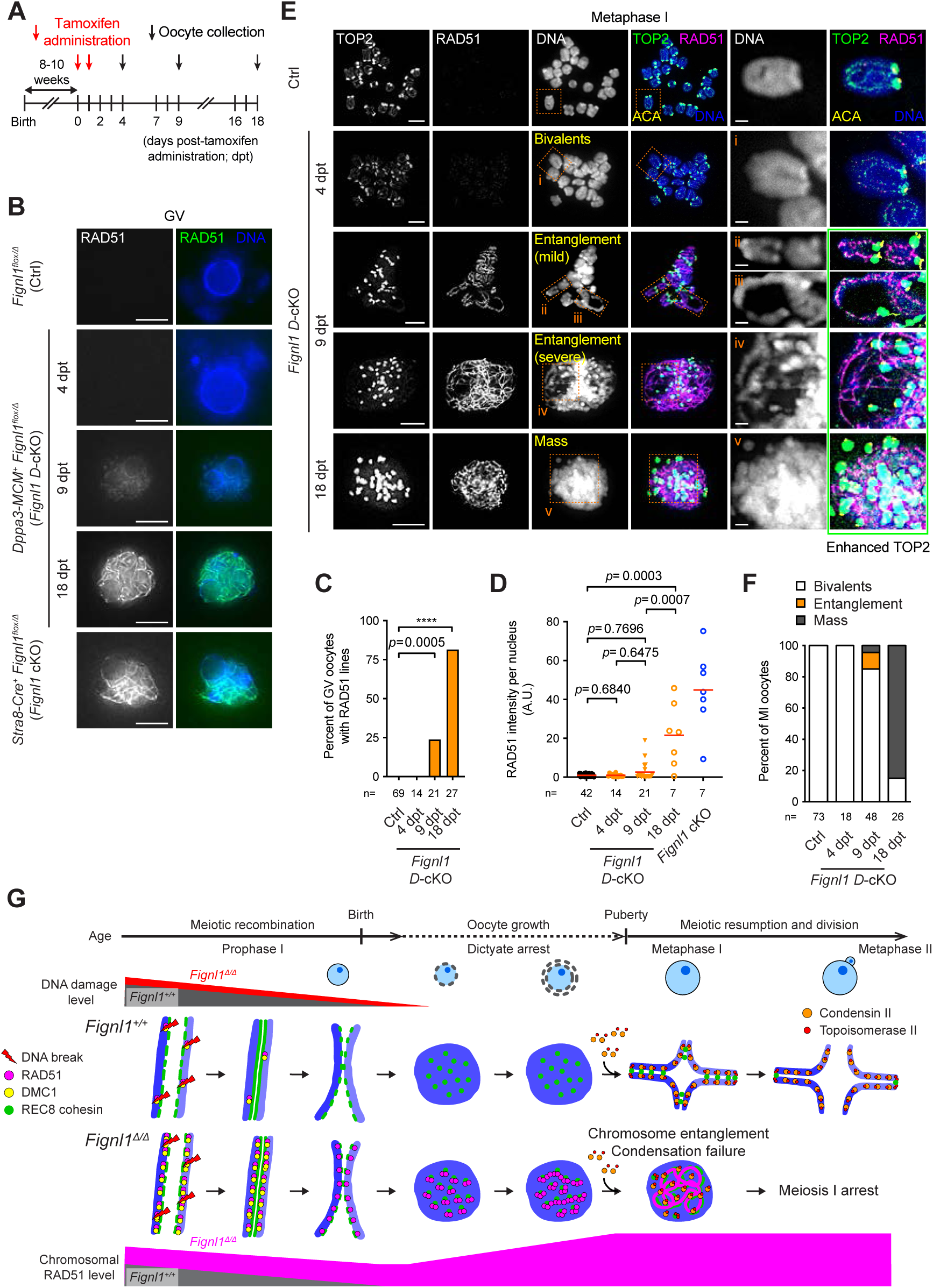
Age-dependent increase in chromosomal RAD51 and the severity of condensation defects by *Fignl1* depletion in growing oocytes. (A) Scheme to deplete *Fignl1* in growing oocytes by *Dppa3-MCM*. (B) Representative images of fixed GV oocytes stained with Hoechst 33342 (DNA, blue) and immunostained for RAD51 (white in the left panels and green in the right panels) from *Fignl1^flox/Δ^* (Ctrl), *Fignl1^flox/Δ^ Dppa3-MCM^+^* (*Fignl1 D*-cKO), and *Fignl1^flox/Δ^ Stra8-Cre^+^* (*Fignl1* cKO) mice at indicated days post-tamoxifen treatment (dpt). Images are optical slices to show the middle of nuclei. (C) Frequencies of GV oocytes with linear immunostaining pattern of RAD51 from Ctrl and *Fignl1 D*-cKO mice at indicated dpt. The results of fisher’s exact tests are indicated in the graph: \*\*\*\**p* ≤ 0.0001. (D) Signal intensities of nuclear RAD51 in fixed GV oocytes from Ctrl (black circles), *Fignl1 D*-cKO (orange filled circles, triangles, and open circles at 4, 9, and 18 dpt, respectively), and *Fignl1* cKO (blue open circles) mice. The red bars are means. The results of two-tailed Mann-Whitney *U*-tests are indicated in the graph. (E) Representative images of chromosome spreads of metaphase-I oocytes stained with DAPI (DNA, white in the left panels and blue in the right panels) and immunostained for TOP2 (white in the left panels and green in the right panels), RAD51 (white in the middle panels and magenta in the right panels), and centromeres (ACA, yellow) from Ctrl and *Fignl1 D*-cKO mice at indicated dpt. Examples of normal bivalents (Ctrl and *Fignl1 D*-cKO at 4 (i) and 9 dpt (ii)), mild entanglement resulting in an abnormally stretched bivalent (*Fignl1 D*-cKO at 9 dpt (iii)), severe entanglement forming a mass-like structure (*Fignl1 D*-cKO at 9 dpt (iv)), and a chromosome mass (*Fignl1 D*-cKO at 18 dpt (v)) are magnified in the right panels. (F) Frequencies of condensation defects in metaphase-I oocytes from Ctrl and *Fignl1 D*-cKO mice at indicated dpt. (G) A summary model of age-dependent RAD51 accumulation and defective meiosis in *Fignl1*-deficient oocytes. RAD51 preferentially accumulates on cohesin REC8-enriched chromosome axes throughout prophase I (before birth) and during the dictyate arrest (after birth). DMC1 also accumulates on chromosome axes during prophase I, but is dissociated by the dictyate stage. Repair of meiotic DSBs is largely efficient and DSBs that remain unrepaired during prophase I are repaired during the dictyate arrest. RAD51 further accumulates during the dictyate arrest as oocytes grow in an age-dependent manner with little increase in DNA damage levels. RAD51 accumulation on chromosome axes impedes proper chromosomal localization of condensin II and topoisomerase II and DNA decatenation by topoisomerase II, leading to massive chromosome entanglement, failure of chromosome condensation and segregation, and meiosis-I arrest. Total numbers of oocytes analyzed are indicated below the graphs. Genotypes of indicated animals are: Ctrl, *Fignl1^flox/Δ^*; *Fignl1 D*-cKO, *Fignl1^flox/Δ^ Dppa3-MCM^+^*; *Fignl1* cKO, *Fignl1^flox/Δ^ Stra8-Cre^+^*. Scale bars in (B) and (E), 10 μm for full nuclei and 2 μm for magnified panels.

Chromosome spreads of metaphase-I oocytes revealed a time-dependent increase in the severity of chromosome condensation defects. While all *Fignl1 D*-cKO oocytes (18/18) at 4 dpt formed normal 20 bivalents like control oocytes, 4.2% (2/48) and 84.6% (22/26) of those at 9 and 18 dpt, respectively, showed a chromosome mass, which was reminiscent of *Stra8-Cre* mediated *Fignl1* cKO oocytes (**Fig. 5E and F, and Fig. S9C**). Notably, 10.4% (5/48) of *Fignl1 D*-cKO oocytes at 9 dpt showed intermediate condensation defects with distinct severity of chromosome entanglement: the mildest had only one abnormally stretched bivalent whereas the most severe oocyte showed massive entanglement, forming a mass-like chromosome structure (**Fig. 5E and F**). All *Fignl1 D*-cKO oocytes with chromosome condensation defects (either entanglement or mass) showed RAD51 accumulation (**Fig. S9D**). These results suggest that cumulative entanglement of multiple chromosomes due to compromised decatenation by TOP2 ultimately results in the formation of the chromosome mass. RAD51 accumulation above a threshold to cause defective chromosome condensation requires a certain time in growing oocytes, thus depends on age. Weak and strong association of RAD51 on chromatid axes of normal bivalent chromosomes in *Fignl1 D*-cKO oocytes at 4 and 9 dpt, respectively (**Fig. 5E, insets i and ii**) further supports the idea that *Fignl1*-deficiency causes preferential RAD51 accumulation on where condensin II and TOP2 bind.

## Discussion

Proper targeting of RAD51 recombinase to DNA is highly regulated in cells for genome stability. In this study, we revealed that *Fignl1* depletion in mouse oocytes leads to aberrant accumulation of RAD51 preferentially on cohesin- and condensin II-enriched chromosome axes and improper localization of condensin II and topoisomerase II on meiotic chromosomes, which results in defective chromosome condensation and arrest at meiosis I. Concomitant increases in chromosomal RAD51 levels and the severity of chromosome condensation defects and chromosome entanglement by *Fignl1* depletion in growing oocytes suggest that accumulation of RAD51 on meiotic chromosomes compromises TOP2 function and causes the defects in chromosome condensation and segregation. Importantly, this promiscuous binding of RAD51 to meiotic chromosomes does not induce obvious DNA damage. We propose that RAD51 accumulation to intact dsDNA, particularly chromosome axes, is a unique type of chromosomal pathology, which interferes with chromosomal processes including condensation and faithful segregation. Oocytes, which are arrested at the dictyate stage for a long period, provides a unique environment for RAD51 to accumulate on meiotic chromosomes in an age-dependent manner (**Fig. 5G**). We infer that aged mammalian oocytes, especially with hypomorphic *FIGNL1* mutations, have higher risk of chromosomal RAD51 accumulation and defective chromosome segregation than young oocytes.

FIGNL1 and FIRRM function as a complex, and their interdependent protein stability leads to nearly identical defects in RAD51 dissociation and DNA repair by their mutations (*3, 6–10, 33, 34*). Nonetheless, little increase in γH2AX levels in *Fignl1-*depleted oocytes, distinct from increased γH2AX levels in *Firrm* cKO oocytes (*21*), suggests that FIRRM alone has residual functions in DNA repair in the absence of FIGNL1 in oocytes. This is consistent with a FIGNL1-independent role of FIRRM in inter-crosslink repair in human somatic cells (*5*). Conversely, stronger RAD51 accumulation in *Fignl1*-depleted oocytes (∼100-200 fold of control oocytes) than *Firrm* cKO oocytes (∼5-fold of the wild-type oocytes) (*21*) suggests that residual FIGNL1 alone could dissociate RAD51, albeit less efficiently, from DNA as it does *in vitro* (*3, 8*).

## Materials and Methods

### Mice

Mice were maintained and euthanized according to the guidelines for the proper conduct of animal experiments (Science Council of Japan) and procedures approved by the Institutional Animal Care and Use Committee at Institute for Protein Research, The University of Osaka (approval ID; 25-02-0) and Department of Biological Sciences, Graduate School of Science, The University of Tokyo (approval ID; A2024S011-01). Mice were housed with food and water provided *ad libitum* and maintained in a temperature-controlled room at 22℃ on a 12h light: 12h dark cycle.

All mice used in this study were congenic to the C57BL/6 background. Mice harboring *Fignl1^flox^* allele were described in our previous study (*3*). *Stra8-Cre* and *Dppa3-MCM* transgenic mice were previously described (*15, 32*) and provided by the RIKEN Animal Resource Center. *Dppa3-MCM*-mediated excision of the *Fignl1^flox^* allele was induced by two consecutive intraperitoneal injections of 0.1 ml of 20 mg/ml Tamoxifen (Sigma, T5648) dissolved in corn oil (Sigma, C8267) with a 24 h interval prior to superovulation.

### Histology

Ovaries were fixed in either 4% paraformaldehyde (Nakarai, 09154-14) or in 10% buffered formalin (Wako, 062-01661) overnight at room temperature, washed twice with PBS, and stored in 70% EtOH at 4°C prior to embedding in paraffin and sectioning. For histological analysis, ovarian paraffin sections of 5 μm thickness were deparaffinized and stained with hematoxylin and eosin (HE&E). For immunostaining, ovarian paraffin sections were deparaffinized and autoclaved in 10 mM sodium citrate buffer (pH 6.0) for 20 min at 121°C for antigen retrieval. Sections were blocked with Blocking One Histo (Nakarai, 06349-64) for 30 min at room temperature followed by incubation with primary and secondary antibodies as described below.

### Collection and culture of oocytes

Germinal-vesicle (GV) stage oocytes were collected from female mice at 4 weeks or 2-3 months superovulated with intraperitoneal injection of 0.1 ml of CARD HyperOva (Kyudo) or 5 IU of pregnant mare serum gonatropin (PMSG) 48 h prior to collection. Surrounding cumulus cells were mechanically removed in M2 medium (Sigma, M7167) containing 0.5 mM N^6^,2’-O-Dibutyryladenosine 3’,5’-cyclic monophosphate (dbcAMP; Sigma, D0627) on a glass/metal heater (Tokai Hit) to maintain 34°C. For *in vitro* maturation, cumulus-free oocytes were washed three times and cultured in pre-equilibrated M16 medium (Sigma, M7292) covered with mineral oil (Sigma, M5310) or liquid paraffin (Nacalai, 26137-85) for 2-3 h, 7 h or 16 h at 37°C under 5% CO_2_ for observation of GV breakdown (GVBD), metaphase I, or metaphase II and polar body exclusion (PBE), respectively.

### Chromosome spreads of fetal oocytes

Surface-spread chromosomes of fetal oocytes were prepared as described previously (*37, 38*) with slight modification. A pair of ovaries was dissected into PBS and incubated in 300 μl of 0.05% Trypsin-EDTA for 10 min at 37°C. After vigorous pipetting to break ovaries into cell suspension, trypsinization was stopped by adding 700 μl of DMEM (Nakarai, 08459-64) containing 10% FBS and Penicillin/Streptomycin (Wako, 168-23191). Cells were pelleted by centrifugation at 200 x*g* for 5 min at room temperature, washed in 200 μl of PBS, and incubated in 20 μl of hypotonic extraction buffer (30 mM Tris-HCl pH 8.0, 50 mM sucrose, 17 mM trisodium citrate dihydrate, 5 mM EDTA, 0.5 mM Dithiothreitol (DTT) and 0.5 mM phenylmethylsulphonyl fluoride (PMSF), pH 8.2-8.3; HEB) for 10 min at room temperature. 20 μl of 100 mM sucrose was added, and 10 μl of cell suspension was placed on a clean glass slide (Matsunami, MAS-01) coated with 50 μl of freshly made 1% paraformaldehyde (PFA) solution pH 9.2 (pH set by 1.25 M sodium borate) containing 0.15% Triton X-100. Slides were placed and dried in a closed humid chamber with hot tap water overnight at room temperature, followed by drying with lids ajar for 3 h and then with lids removed for 1-2 hr. Slides were washed once for 5 min in deionized water and twice for 5 min in 0.4% DryWell (Fujifilm) in a coplin jar and air-dried at room temperature. Slides were either directly processed for immunostaining or stored in aluminum foil at -80°C prior to immunostaining.

### Chromosome spreads of metaphase-I oocytes

Chromosome spreads of metaphase-I oocytes were prepared as described previously (*38*) with slight modification. Cultured oocyte at 7 h from dbcAMP washout were transferred to M2 medium then Tyrode’s solution (Sigma, T1788) to remove zona pellucida (ZP). ZP-free oocytes were washed once in M2 medium and transferred to 5 μl of 1% PFA solution pH 9.2 (pH set by 1.25 M sodium borate) containing 0.15% Triton X-100 and 3 mM DTT placed in a well of a 12-well glass slide (Matsunami, TF1205M). Slides were air-dried overnight at room temperature and either directly processed for immunostaining or stored in aluminum foil at 4°C prior to immunostaining.

### Immunofluorescence staining

Oocytes were permeabilized in PBS containing 0.5% Triton X-100 for 1 min at room temperature followed by fixation in 4% PFA (Nakarai, 09154-14) for 20 min at room temperature, washed once and stored in PBS at 4°C prior to immunostaining. For observation of PBE, oocytes were directly fixed in 4% PFA for 20 min at room temperature followed by permeabilization in PBS containing 0.2% Triton X-100 for 20 min. Fixed and permeabilized oocytes were blocked with blocking buffer (0.33% normal goat serum, 1% bovine serum albumin (BSA), 1x TBS pH 8.0, 0.017% Triton X-100, 0.017 % sodium azide) for 1 h at room temperature and incubated with primary antibodies in antibody dilution buffer (10% normal goat or donkey serum, 3% BSA, 1x TBS pH 8.0, 0.05% of Triton X-100, 0.05% sodium azide) overnight at 4°C. Oocytes were washed once with PBS for 5 min, incubated with secondary antibodies in antibody dilution buffer for 1 h at room temperature, and washed once with PBS for 5 min prior to imaging. Immunostaining of chromosome spreads was performed as previously described (*3*). Primary antibodies used are: mouse anti-RAD51 (1:100; Novus, NB100-148), rat anti-α-tubulin (1:500; Santa Cruz, sc-53029), mouse anti-γH2AX (1:500; Millipore, 05-636), rabbit anti-NCAPD3 (1:500; gift from T. Hirano), rabbit anti-REC8 (1:500; gift from T. Kitajima), rabbit anti-TOP2 (1:400; Abcam, ab109524), mouse anti-TOP2A (1:100; Santa Cruz, sc-365916), rabbit anti-H3K9me3 (1:500; Abcam, ab8898), rabbit anti-DDX4 (1:500; Abcam, ab13840), mouse anti-SYCP3 (1:750; Abcam, ab97672), rat anti-SYCP3 (1:5,000; in house (*3*)), rabbit anti-DMC1 (1:200; Santa Cruz, sc-22768), rabbit ant-RPA2 (1:200; Abcam, ab76420), rat anti-RPA2 (1:100; Cell Signaling Technology, 2208), mouse anti-MLH1 (1:30; Cell Signaling Technology, 3515), human anti-centromere (1:200; Antibodies Incorporated, 15-235). Secondary antibodies used are: Alexa Fluor 488-conjugated goat anti-mouse (1:1,000; Invitrogen, A-11029) or anti-rat (1:1,000; Invitrogen, A-11006) or anti-rabbit (1:1,000; Invitrogen, A-11034), Alexa Fluor 594-conjugated goat anti-rat (1:1,000; Invitrogen, A-11007) or anti-human (1:1,000; Invitrogen, A-11014), Alexa Fluor 647-conjugated goat anti-rabbit (1:500; Invitrogen, A-21245) or anti-mouse (1:500; Invitrogen, A-21235).

### Image acquisition

Images of immunostained fixed oocytes, chromosome spreads and ovary sections were acquired using a computer-assisted fluorescence microscope system (DeltaVision; Applied Precision) equipped with a 60× NA 1.4 oil immersion objective and a 10x NA 0.4 objective. Whole oocyte images were acquired as *z* stacks at 0.5 μm intervals. Image deconvolution was performed using an image workstation (softWoRx version 6.5.2; Applied Precision) and afterward processed using Imaris (version 9.2.1; Oxford Instruments) and Photoshop (version 23.2.2; Adobe) software tools. Images of HE&E-stained ovary sections were acquired using a EVOS XL Core Cell Imaging system (Life technologies) and processed using Photoshop software.

### Live imaging

GV stage oocytes were collected as described above. *In vitro*–transcribed mRNAs were microinjected into GV oocytes maintained in M2 medium supplemented with 150 or 500 μM dbcAMP and overlaid with liquid paraffin to prevent evaporation, using a Piezo-driven micromanipulator (Prime Tech). After injection, oocytes were cultured in M2 medium containing 150 or 500 μM dbcAMP for ∼2 h at 37°C under 5% CO_2_. For live imaging, oocytes were transferred to drops of pre-equilibrated M16 medium covered with liquid paraffin in glass-bottom dishes (Matsunami, D11130H) and maintained at 37°C under 5% CO_2_ during observation. Imaging was performed on an Olympus IX71 inverted microscope equipped with a Yokogawa CSU-X1 spinning-disk confocal scanner, a UPlanSApo 60× silicone-oil immersion objective (Olympus), and an Andor iXon DU-897E-CS0-#BV EMCCD camera, controlled by MetaMorph software. Excitation was provided by 488 nm and 568 nm lasers. *Z*-stacks consisted of one transmitted-light (bright-field) optical section and 21 fluorescence sections acquired at 4-µm intervals, with time-lapse imaging performed at 5- or 20-min intervals.

Capped mRNAs were synthesized *in vitro* from enzymatically linearized plasmids using the T7 RiboMAX Large Scale RNA Production System (Promega) with Ribo m⁷G Cap Analog (Promega). Expression constructs were kindly provided as follows: a T7 promoter/poly(A)-containing backbone plasmid from K. Yamagata (Kindai University), the histone H2B construct from H. Kimura (Institute of Science Tokyo), and mouse MAP4 cDNA cloned from testis. The MTBD of MAP4 was defined as spanning from Proline 659 to the C-terminal residue.

### TRIM-Away

Capped HaloTag–TRIM21 mRNA and antibodies were co-injected into GV oocytes using a Piezo-driven micromanipulator as described above. The antibodies used were anti-REC8 (0.164-0.636 mg/ml, gift from T. Kitajima), anti-RAD51 (0.07-0.088 mg/ml, Novus, NB100-148), and normal IgG (0.161-0.429 mg/ml, MBL, PM035) as a negative control. Immediately after injection, oocytes were transferred to pre-equilibrated M16 medium covered with liquid paraffin and cultured at 37°C under 5% CO_2_ for 7 h, after which chromosome spreads were prepared as described above. TRIM21 cDNA, cloned from NIH-3T3 cells, was fused in-frame downstream of a HaloTag sequence via a 9-amino acid linker and inserted into the T7 promoter/poly(A)-containing backbone vector. Capped mRNA was synthesized *in vitro* from enzymatically linearized plasmids using the T7 RiboMAX Large Scale RNA Production System (Promega) with Ribo m⁷G Cap Analog, as described above.

### Image analysis

Single nuclei were manually cropped, and numbers of foci were counted by the auto-thresholding signal intensity in Imaris software (version 9.2.1; Oxford Instruments). Signal intensities on chromosomes or in nuclei were measured from non-deconvolved images with matched exposure using ImageJ software (Fiji version 2.9.0) by subtracting signal intensities on the DAPI or Hoechst 33342 staining from those of regions of interest manually drawn next to chromosomes (chromosome spreads), nuclei but inside cytoplasm (fixed oocytes), or sections (ovary sections) as backgrounds. Signal intensities of RAD51 and γH2AX in fixed oocytes were quantified in an optical slice in the middle of the nucleus and by integrating signal intensities of each optical slice containing the nucleus, respectively. Chromosome volume was measured by a 3D surface rendering with mRFP-H2B signals in Imaris software.

Meiotic prophase I stages were defined by SYCP3 and SYCP1 staining using standard criteria. Leptonema was defined by short SYCP3 stretches without evidence of synapsis determined by SYCP1 staining from E15.5 ovaries. Zygonema was defined by longer stretches of SYCP3 with various degrees of synapsis from E15.5 ovaries. Pachynema was defined by the full synapsis of all chromosomes from E18.5 ovaries. Diplonema was defined by de-synapsis with various degrees of residual synapsis from 1 dpp ovaries. Dictyate was defined by short, ragged SYCP3 stretches without evidence of synapsis from 1 dpp ovaries.

For ovary sections, numbers of follicles containing an oocyte with a clearly visible nucleus were manually counted for every fifth section from one ovary per animal. The cumulative counts were multiplied by a correction factor of 5 to estimate the total number of oocytes per ovary as described previously (*39*).

### Chromatin immunoprecipitation – sequencing (ChIP-seq)

ChIP was performed as described previously (*3*). Testes from male mice at 13-15 dpp were dissected, the tunica albuginea was removed, and seminiferous tubules were crosslinked in 5 or 10 ml of 1% formaldehyde (Pierce, 28908) in PBS (Nakarai, 14249-24) for 10 min at room temperature. Crosslinking was quenched by adding 2.5 M glycine to a final concentration of 125 mM. After 5 min of incubation at room temperature, cells were washed once with 10 ml of cold PBS, and the cell pellet was snap-frozen on liquid nitrogen and stored at -80℃. The frozen cell pellet from 4-5 pairs of testes was pooled and suspended in 500 μl of cell lysis buffer (0.25% Triton X-100, 10 mM EDTA, 0.5 mM EGTA, 10 mM Tris-HCl pH 8.0) supplemented with protease inhibitor (Roche, 04693159001), incubated for 10 min on ice, and centrifuged at 300 x *g* for 3 min at 4℃. The nuclei pellet was washed in 500 μl of lysis wash buffer (200 mM NaCl, 1 mM EDTA, 0.5 mM EGTA, 10 mM Tris-HCl pH 8.0) supplemented with protease inhibitor, resuspended in 80 μl of RIPA buffer (10 mM Tris-HCl pH 8.0, 1 mM EDTA, 0.5 mM EGTA, 1% Triton X-100, 0.1% sodium deoxycholate, 0.1% SDS, 140 mM NaCl) supplemented with protease inhibitor, and sonicated on the Covaris M220 for 5 min 30 sec at 5-9°C with a setting of Peak Power 50.0, Duty Factor 20.0, and Cycles/Burst 200. After adding 370 μl of RIPA buffer supplemented with protease inhibitor, sheared nuclei were centrifuged at 20,400 x *g* for 15 min at 4℃ to remove insoluble debris. 400 μl of the chromatin-containing supernatant was mixed with 50 μl of Dynabeads protein G bound to 10 μg of RAD51 antibody (Novus, NB100-148) and incubated overnight on a rotating wheel at 4℃ for immunoprecipitation. Beads were washed once with 1 ml of cold ChIP wash buffer 1 (0.1% SDS, 1% Triton X-100, 2 mM EDTA, 20 mM Tris-HCl pH 8.0, 150 mM NaCl), 1 ml of cold ChIP wash buffer 2 (0.1% SDS, 1% Triton X-100, 2 mM EDTA, 20 mM Tris-HCl pH 8.0, 500 mM NaCl), 1 ml of cold ChIP wash buffer 3 (0.25 M LiCl, 1% NP-40, 1% sodium deoxycholate, 1 mM EDTA, 10 mM Tris-HCl pH 8.0), and twice with 1 ml of cold TE (1 mM EDTA and 10 mM Tris-HCl pH 8.0), followed by elution with 100 μl of elution buffer (0.1 M NaHCO_3_ and 1% SDS) for 30 min at 65℃ with vortexing every 10 min. The elutes were incubated overnight at 65℃ for de-crosslinking, neutralized by adding 4 μl of 1M Tris-HCl (pH 6.5) and 2 μl of 0.5 M EDTA, and RNA was digested by adding 2 μl of 10 mg/ml RNase-A and incubation for 15 min at 37°C. After adding 5 μl of 10 mg/ml Proteinase-K (Wako, 165-21043) and incubation for 1 h at 55℃, DNA was purified using QIAquick PCR Purification Kit (QIAGEN).

IP DNA was further sonicated on the Covaris M220 for 8 min 30 sec at room temperature with a setting of Peak Power 75.0, Duty Factor 10.0, and Cycles/Burst 200. Sequencing libraries were prepared from ∼10 ng of IP DNA using NEBNext Ultra II Library Prep Kit for Illumina (NEB, E7645), NEBNext Multiplex Oligos for Illumina (NEB, E6440), and AMPureXP beads (Beckman Coulter, A63880) according to manufacturer’s instructions and sequenced on NovaSeq X Plus in a 150 bp paired-end run. Around 70 million paired reads were generated per sample. Resulting reads were processed using fastp (*40*) and mapped to the mm10 genome using bowtie2 (*41*) with default settings. Mapped reads from two independent samples of each genotype were averaged and a bigwig file per genotype was generated using Deeptools (*42*).

### *In vitro* DNA relaxation assay

Human RAD51 was purified as described previously (*43*). Supercoiled pBluescript II SK+ (2,961 bp) was purified using DNA extraction kit (QIAGEN). Indicated concentrations of human RAD51 was incubated with 1.28 nM (3.8 μM in bp) of supercoiled pBluescript II SK+ plasmid at 37 °C for 5 min prior to the addition of human TOP2A (5 or 2.5 units, Topogen, TG2000H) in a reaction mixture containing 1x Topo II assay buffer (Topogen). The reactions were incubated at 37°C for 20 min. When indicated, BamHI (15 units, TaKaRa) was added and the reaction was further incubated for an additional 5 min. The reactions were terminated by deproteination with 0.45% SDS and 0.45 mg/ml Proteinase K followed by the incubation at 37°C for 10 min. The products were analyzed by electrophoresis on 1% agarose gel in TAE buffer and visualized by SYBR Gold staining (Thermo Fisher Scientific). DNA was detected by LAS4000 (GE healthcare).

## Statistical analysis and reproducibility

Statistical analyses were performed using Graphpad Prism software version 9.5.1 and R version 4.2.2 (https://www.r-project.org). Bars in figures and values in manuscripts indicate means otherwise mentioned. Statistic parameters and tests and sample sizes are described in the figures and/or figure legends. Sample sizes were not predetermined using any statistical tests.

At least two animals of each genotype were analyzed and similar results were obtained otherwise mentioned. ≥2 independent experiments/animals were pooled for quantification. Two independent experiments were performed for biochemical analysis and similar results were obtained.

## Acknowledgments

We thank Drs. T. Hirano and T. Kitajima for antibodies; Dr. M. Schuh for TRIM-Away constructs; Drs. T. Kitajima and O. Takenouchi for advice; Ms. S. Aoyama, A. Maeda, and M. Yasumura for mouse care and technical supports; and the Shinohara lab and Ohsugi lab for supports and discussions. This work was supported by JSPS KAKENHI Grants 20K15716 and 24K09545 (to M.I.), 22K06218 and 25K09610 (to T.K.), 23K20059 (to A.F.), and 23K18091 and 23K27178 (to MO), grants from Takeda Science Foundation (to M.I. and A.S.), Japan Health Foundation and Mochida Memorial Foundation (to M.I.), The Naito Foundation (to A.F.), and The Uehara Memorial Foundation, Daiichi Sankyo Foundation of Life Science, The NOVARTIS Foundation Japan for the Promotion of Science, and The Institute for Fermentation, Osaka (to A.S.). M.I., S.S., and A.F. were supported by the new field development support program funded by Institute for Protein Research, the University of Osaka.

## Author contributions

M.I., S.S., and A.S. conceived the study and designed the experiments. M.I., S.S., and R.O. performed the experiments and analyzed the data. T.K. and M.K. performed live imaging and TRIM-Away experiments. A.F. performed biochemical experiments. M.I. wrote the manuscript with input and edits from all authors.

## Competing interest

Authors declare that they have no competing interests.

## Notes

### Competing Interest Statement

The authors have declared no competing interest.

