## Supplemental Figures for "Age-dependent accumulation of RAD51 on non-damaged chromosomes prevents chromosome segregation in mammalian oocytes"

Ito et al., Figure S1

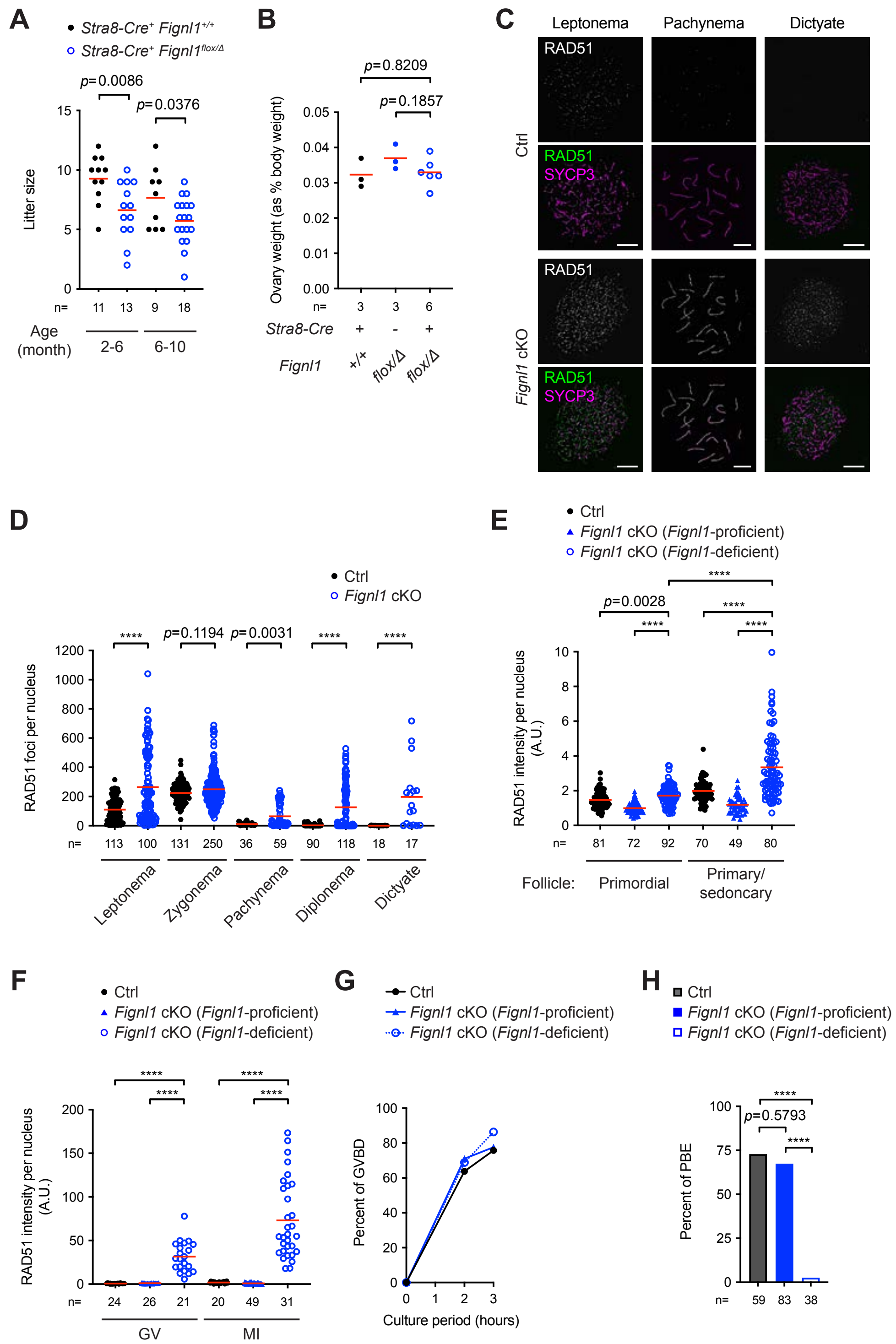


**Fig. S1. Defective oogenesis with aberrant RAD51 accumulation in *Fignl1* cKO mice.**

(A) Litter size of *Fignl1*$^{+/+}$ *Stra8-Cre*$^{+}$ (Ctrl) and *Fignl1*$^{flox/\Delta}$ *Stra8-Cre*$^{+}$ (*Fignl1* cKO) female mice at indicated ages bred to wild-type male mice. The red bars are means. The results of two-tailed unpaired *t*-tests are indicated in the graph. Numbers of animals analyzed are indicated below the graph.

(B) Ovary weights of adult mice (8-10 weeks) with indicated genotypes. The red bars are means. The results of two-tailed unpaired *t*-tests are indicated in the graph. Numbers of animals analyzed are indicated below the graph.

(C) Representative images of oocyte chromosome spreads immunostained for RAD51 (white in the top panels and green in the bottom panels) and SYCP3 (magenta) at indicated meiotic prophase-I stages from Ctrl and *Fignl1* cKO mice. Oocytes at leptotene, pachytene and dictyate stages were from ovaries at embryonic days 15.5 (E15.5), 18.5 (E18.5), and 1 day post-partum (dpp), respectively. Scale bars, 10 μm.

(D) Focus counts of RAD51 in fetal oocytes at different meiotic prophase-I stages from Ctrl (black circles) and *Fignl1* cKO (blue open circles) mice. Oocytes at leptotene, zygotene, pachytene, diplotene, and dictyate stages were from ovaries at E15.5, E15.5, E18.5, 1 dpp, and 1 dpp, respectively. The red bars are means. The results of two-tailed Mann-Whitney *U*-tests are indicated in the graph. Total numbers of oocytes analyzed are indicated below the graph.

(E) Signal intensities of nuclear RAD51 in Ctrl (black circles), *Fignl1*-proficient (blue triangles), and *Fignl1*-deficient (blue open circles) oocytes in ovary sections from 18-dpp animals. The red bars are means. The results of two-tailed Mann-Whitney *U*-tests are indicated in the graph: ****$p \leq 0.0001$. Total numbers of oocytes analyzed are indicated below the graph.

(F) Signal intensities of nuclear RAD51 in fixed Ctrl (black circles), *Fignl1*-proficient (blue triangles), and *Fignl1*-deficient (blue open circles) oocytes at GV and metaphase-I stages. The red bars are means. The results of two-tailed Mann-Whitney *U*-tests are indicated in the graph. Total numbers of oocytes analyzed are indicated below the graph.

(G) Timing and efficiency of germinal-vesicle break down (GBVD) in Ctrl (black circles and line), *Fignl1*-proficient (blue triangles and line), and *Fignl1*-deficient (blue open circles and dashed line) oocytes. Total numbers of oocytes analyzed from five animals of each genotype are: 69 and 62 in Ctrl; 62 and 76 in *Fignl1*-proficient; 45 and 44 in *Fignl1*-deficient oocytes at 2 h and 3 h, respectively.

(H) Efficiency of polar body extrusion (PBE), an indicative of meiosis I division, in Ctrl (a gray bar), *Fignl1*-proficient (a blue bar), and *Fignl1*-deficient (a blue open bar) oocytes after 16 h of culture. The results of Fisher's exact tests are indicated in the graph: ****$p \leq 0.0001$. Total numbers of oocytes analyzed from two animals of each genotype are indicated below the graph.

Genotypes of indicated animals are: Ctrl, *Fignl1*$^{+/+}$ *Stra8-Cre*$^{+}$ in (A) and (E) and *Fignl1*$^{flox/+}$ *Stra8-Cre*$^{+}$ in (C), (D), (F), (G), and (H); *Fignl1* cKO, *Fignl1*$^{flox/\Delta}$ *Stra8-Cre*$^{+}$. In (E)-(H), RAD51-negative and -positive oocytes from *Fignl1* cKO mice were separately analyzed as *Fignl1*-proficient and -deficient oocytes, respectively.

Ito et al., Figure S2

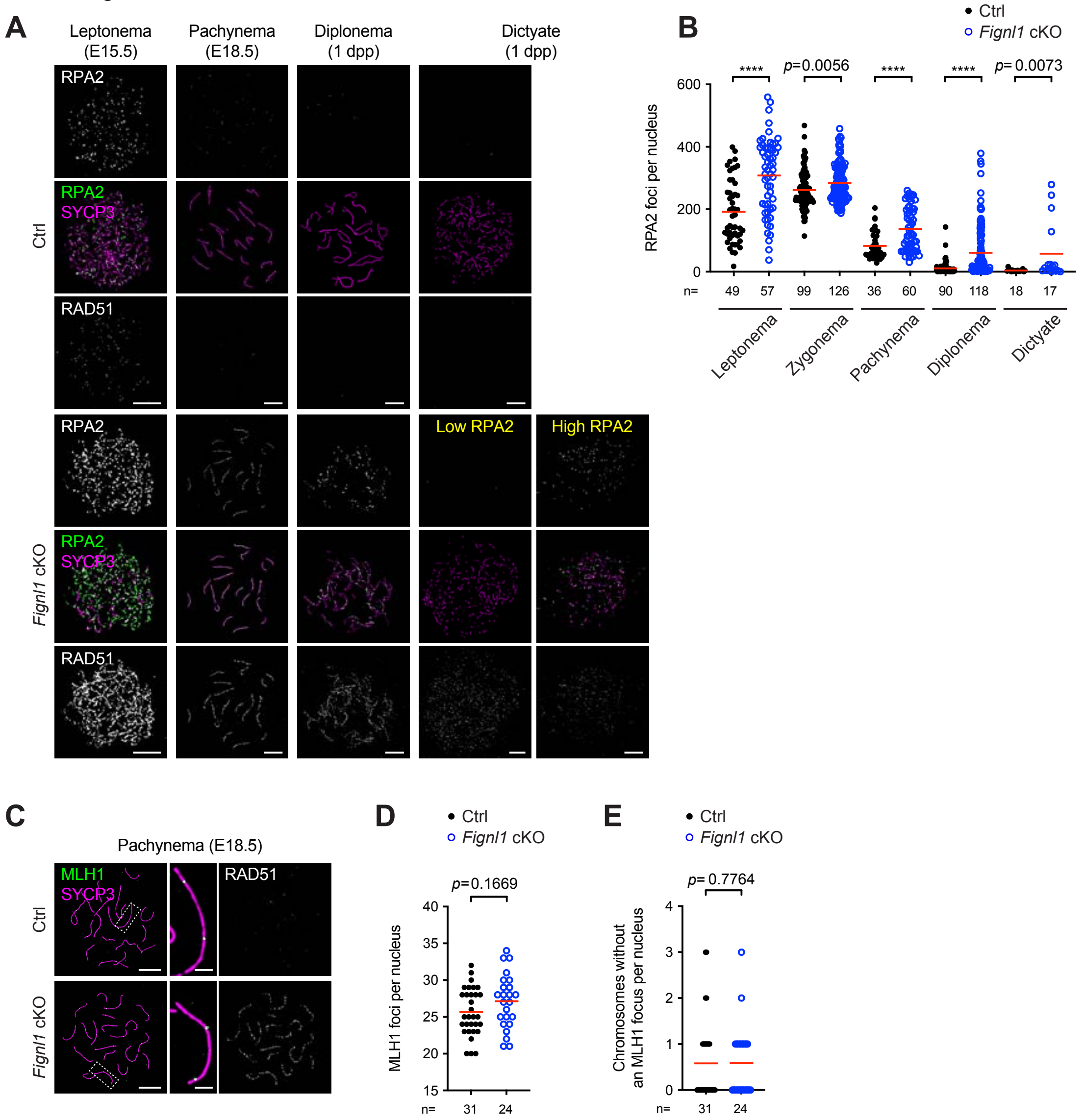


**Fig. S2. Largely normal repair of meiotic DSBs and crossover formation during prophase I in *Fignl1*-deficient oocytes.**

(A) Representative images of oocyte chromosome spreads immunostained for RPA2 (white in the top panels and green in the middle panels), SYCP3 (magenta), and RAD51 (white in the bottom panels) at indicated meiotic prophase I stages from Ctrl and *Fignl1* cKO mice. For dictyate oocytes from *Fignl1* cKO mice, nuclei with high and low RPA2 focus counts are shown.

(B) Focus counts of RPA2 in fetal oocytes at different meiotic prophase-I stages from Ctrl (black circles) and *Fignl1* cKO (blue open circles) mice. The red bars are means.

(C) Representative images of chromosome spreads of pachytene oocytes immunostained for MLH1 (green), SYCP3 (magenta), and RAD51 (white) from Ctrl and *Fignl1* cKO mice.
(D) Focus counts of MLH1 in pachytene Ctrl (black circles) and *Fignl1* cKO (blue open circles) oocytes. The red bars are means.
(E) Numbers of chromosomes that lack an MLH1 focus in pachytene Ctrl (black circles) and *Fignl1* cKO (blue open circles) oocytes. The red bars are means.
Oocytes at leptotene, zygotene, pachytene, diplotene, and dictyate stages were from ovaries at E15.5, E15.5, E18.5, 1 dpp, and 1 dpp, respectively. Pachytene oocytes with ≥100 RAD51 foci in *Fignl1* cKO mice were selectively analyzed in (D) and (E). The results of two-tailed Mann-Whitney *U*-tests are indicated in the graphs: $****p \leq 0.0001$. Total numbers of oocytes analyzed are indicated below the graphs. Genotypes of indicated animals are: Ctrl, $Fignl1^{flox/+}$ *Stra8-Cre*$^{+}$; *Fignl1* cKO, $Fignl1^{flox/\Delta}$ *Stra8-Cre*$^{+}$. Scale bars in (A) and (C), 10 μm for full nuclei and 2 μm for magnified panels.

Ito et al., Figure S3

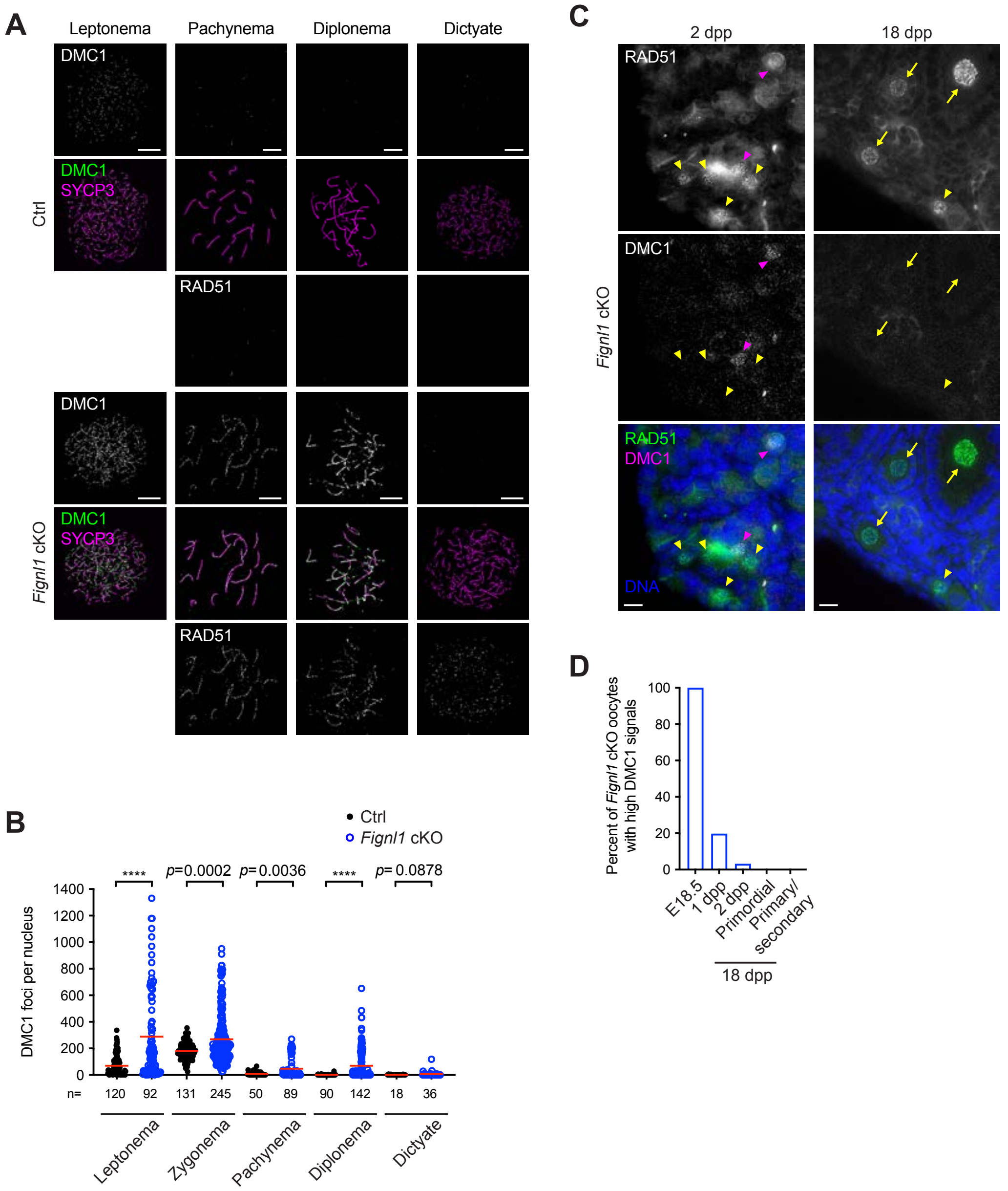


**Fig. S3. Altered DMC1 dynamics throughout meiotic prophase I and the dictyate stage in *Fignl1*-deficient oocytes.**

(A) Representative images of oocyte chromosome spreads immunostained for DMC1 (white in the top panels and green in the middle panels), SYCP3 (magenta), and RAD51 (white in the bottom panels) at indicated meiotic prophase I stages from Ctrl and *Fignl1* cKO mice.

(B) Focus counts of DMC1 in fetal oocytes at different meiotic prophase I stages from Ctrl (black circles) and *Fignl1* cKO (blue open circles) mice. The red bars are means. The results of two-tailed Mann-Whitney *U*-tests are indicated in the graphs: ****$p \leq 0.0001$. Total numbers of oocytes analyzed are indicated below the graphs.

(C) Representative images of ovary sections stained with DAPI (DNA, blue) and immunostained for RAD51 (white in the top panels and green in the bottom panels) and DMC1 (white in the middle panels and magenta in the bottom panels) from *Fignl1* cKO mice at 2 and 18 dpp. Magenta and yellow arrowheads indicate primordial follicles containing *Fignl1* cKO oocytes with high and low DMC1 signals, respectively. Yellow arrows indicate primary and secondary follicles containing *Fignl1* cKO oocytes with low DMC1 signals.

(D) Frequencies of *Fignl1* cKO oocytes with high DMC1 signals at indicated ages and follicular stages. DMC1 signals were analyzed by oocyte chromosome spreads for pachytene oocytes at E18.5 and ovary sections at 1, 2, and 18 dpp. Total numbers of oocytes analyzed are: 13 from one animal at E18.5, 111 from one animal at 1 dpp; 186 from two animals at 2 dpp; 118 in primordial and 167 in primary/secondary follicles from two animals at 18 dpp.

Oocytes at leptotene, zygotene, pachytene, diplotene, and dictyate stages were from ovaries at E15.5, E15.5, E18.5, 1 dpp, and 1 dpp, respectively. Genotypes of indicated animals are: Ctrl, *Fignl1*$^{flox/+}$ *Stra8-Cre*$^{+}$; *Fignl1* cKO, *Fignl1*$^{flox/\Delta}$ *Stra8-Cre*$^{+}$. Scale bars in (A) and (C), 10 μm.

Ito et al., Figure S4

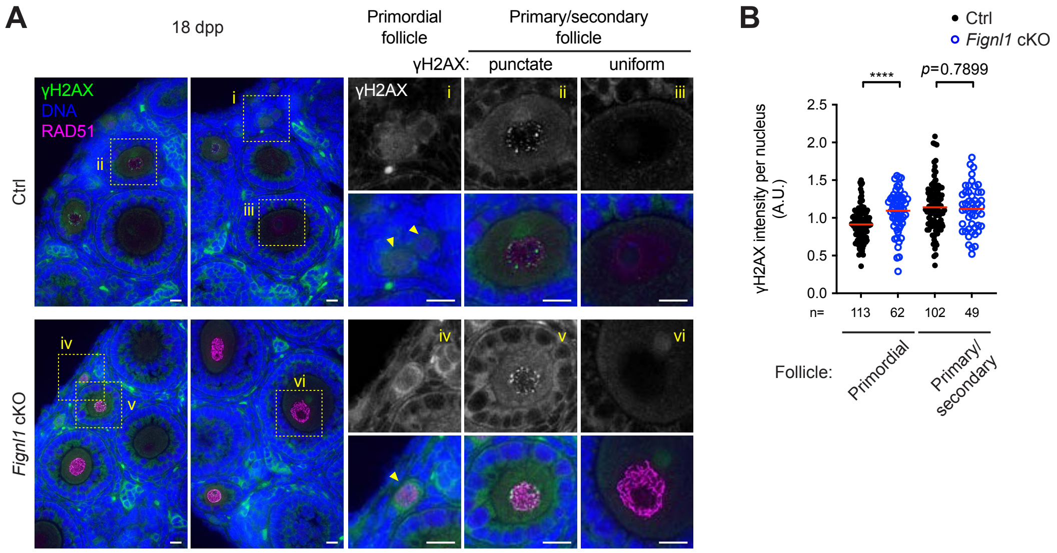


**Fig. S4. Efficient DNA repair in *Fignl1*-deficient dictyate oocytes.**

(A) Representative images of ovary sections stained with DAPI (DNA, blue) and immunostained for γH2AX (white and green) and RAD51 (magenta) from Ctrl and *Fignl1* cKO mice at 18 dpp. Examples of primordial (i and iv) and primary/secondary follicles with (ii and v) and without (iii and vi) punctate γH2AX signals are magnified in the right panels. Scale bars, 10 μm

(B) Signal intensities of nuclear γH2AX in Ctrl (black circles) and *Fignl1* cKO (blue open circles) oocytes in ovary sections from 18-dpp animals. The red bars are means. The results of two-tailed Mann-Whitney *U*-tests are indicated in the graphs: $^{****}p \leq 0.0001$. Total numbers of oocytes analyzed are indicated below the graphs.

Genotypes of indicated animals are: Ctrl, $Fignl1^{+/+}$ $Stra8\text{-}Cre^{+}$; *Fignl1* cKO, $Fignl1^{flox/\Delta}$ $Stra8\text{-}Cre^{+}$.

Ito et al., Figure S5

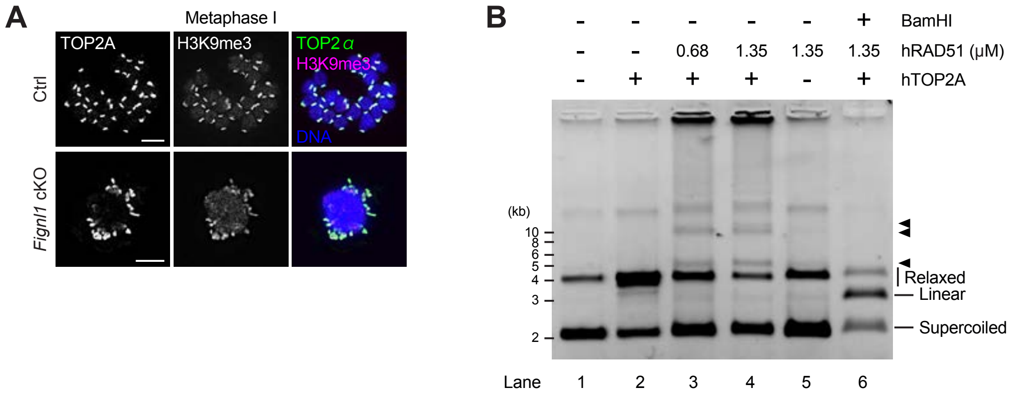


**Fig. S5. Improper chromosomal localization and compromised function of topoisomerase II on RAD51-bound DNA.**

(A) Representative images of chromosome spreads of metaphase-I oocytes stained with DAPI (DNA, blue) and immunostained for TOP2A (white in the left panels and green in the right panels) and trimethylation of histone H3 at lysine 9 (H3K9me3, white in the middle panels and magenta in the right panels) from Ctrl (*Fignl1*$^{flox/+}$ *Stra8-Cre*$^{+}$) and *Fignl1* cKO (*Fignl1*$^{flox/\Delta}$ *Stra8-Cre*$^{+}$) mice. Scale bars, 10 μm.

(B) A representative gel image of DNA relaxation assay. Reaction conditions were as described for Fig. 3F but with 2.5 units of TOP2A instead of 5 units. Migration positions of supercoiled, linear, and relaxed or open circular DNA are indicated. Black arrowheads indicate slower migration products specifically detected or enriched in the presence of RAD51 in the TOP2A reaction (lanes 3 and 4).

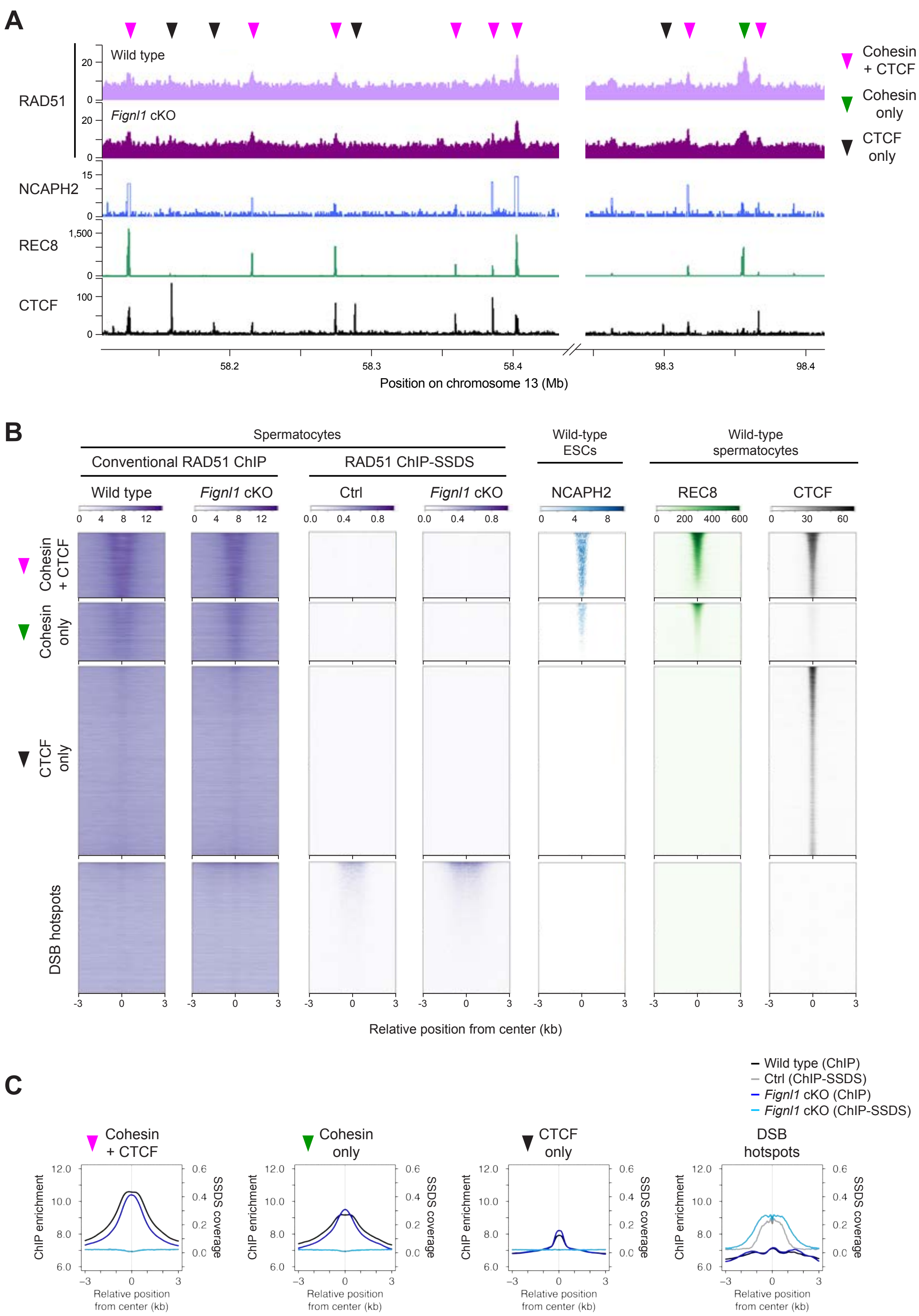


**Fig. S6. Preferential RAD51 binding to intact dsDNA on chromosome axes.**
(A) ChIP signals of RAD51 in wild-type (light purple) and *Fignl1* cKO (dark purple)

spermatocytes with NCAPH2 in ES cells (blue) and REC8 (green) and CTCF (black) in wild-type spermatocytes at indicated regions on mouse chromosome 13. Magenta, Green, and black triangles indicates cohesin + CTCF, cohesin only, and CTCF only sites, respectively. ChIP data for NCAPH2, REC8, and CTCF are from previous studies (*28, 36*).

(B) Heatmaps of conventional ChIP and ChIP-SSDS signals of RAD51 in spermatocytes (purple) around chromosome axes and DSB hotspots. Chromosome axis sites are ordered as in Fig. 4A and 13,944 DSB hotspots are ordered by SPO11-oligo counts (*44*). Heatmaps of *Fignl1* cKO (conventional ChIP), NCAPH2, REC8, and CTCF around chromosome axis sites are reproduced from Fig. 4A. RAD51 ChIP-SSDS data are from our previous study (*3*).

(C) Metaplots of ChIP and ChIP-SSDS signals of RAD51 around chromosome axis sites and DSB hotspots. Smoothed ChIP and ChIP-SSDS signals of RAD51 in wild-type or Ctrl (black and gray, respectively) and *Fignl1* cKO (blue and light blue, respectively) spermatocytes around ±3 kb of cohesin + CTCF sites, cohesin only sites, CTCF only sites, and DSB hotspots used for heatmap representation are shown.

Genotypes of indicated animals are: Ctrl, *Fignl1*$^{+/+}$ *Stra8-Cre*$^{+}$; *Fignl1* cKO, *Fignl1*$^{flox/\Delta}$ *Stra8-Cre*$^{+}$.

Ito et al., Figure S7

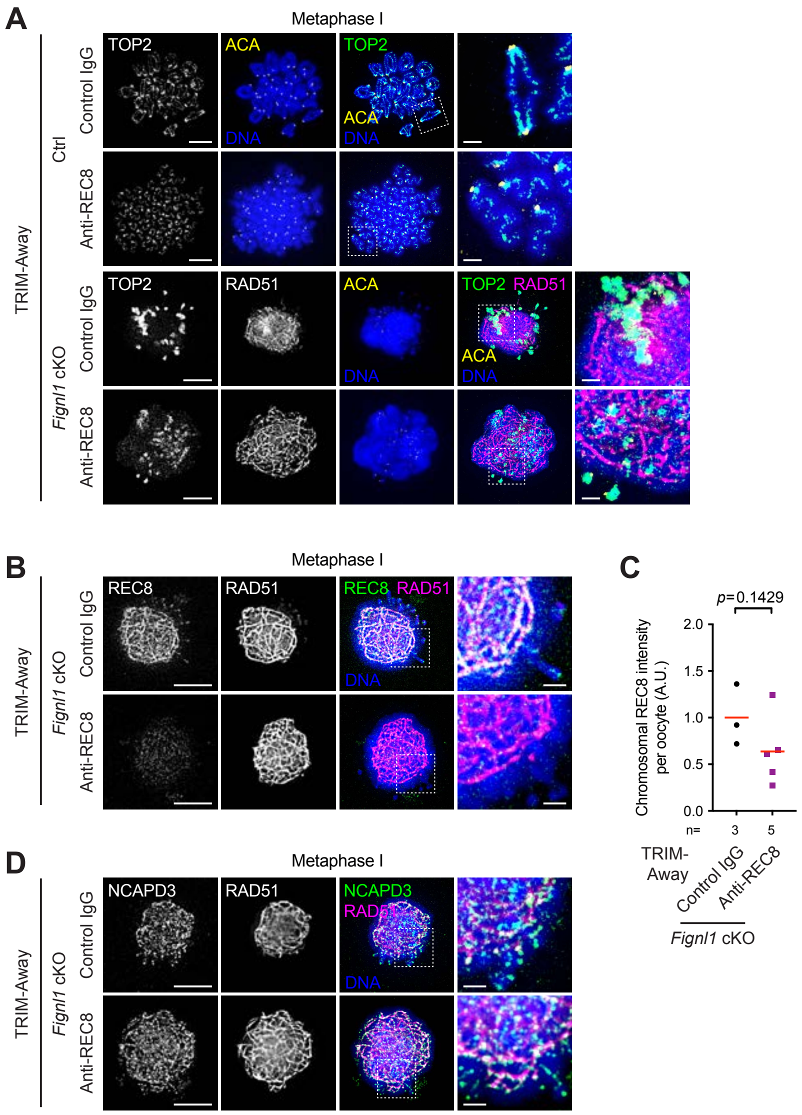


**Fig. S7. Little effects of REC8 depletion on defective chromosome condensation in metaphase-I *Fignl1*-deficient oocytes.**

(A) Representative images of chromosome spreads of control and REC8 TRIM-Away oocytes at metaphase I stained with DAPI (DNA, blue) and immunostained for TOP2 (white in the left panels and green in the right panels), RAD51 (white in the left panels and magenta in the right panels), and centromeres (ACA, yellow) from Ctrl and *Fignl1* cKO mice.

(B) Representative images of chromosome spreads of control and REC8 TRIM-Away oocytes at metaphase I stained with DAPI (DNA, blue) and immunostained for REC8 (white in the left panels and green in the right panels) and RAD51 (white in the left panels and magenta in the right panels) from *Fignl1* cKO mice.

(C) Signal intensities of chromosomal REC8 on chromosome spreads of control (black circles) and REC8 (purple squares) TRIM-Away oocytes at metaphase I from *Fignl1* cKO mice. The red bars are means. The result of two-tailed Mann-Whitney *U*-test is indicated in the graph. Total numbers of oocytes analyzed are indicated below the graph.
(D) Representative images of chromosome spreads of control and REC8 TRIM-Away oocytes at metaphase I stained with DAPI (DNA, blue) and immunostained for NCAPD3 (white in the left panels and green in the right panels) and RAD51 (white in the left panels and magenta in the right panels) from *Fignl1* cKO mice.
Genotypes of indicated animals are: Ctrl, *Fignl1*$^{flox/+}$ *Stra8-Cre*$^{+}$; *Fignl1* cKO, *Fignl1*$^{flox/\Delta}$ *Stra8-Cre*$^{+}$. Scale bars in (A), (B), and (D), 10 μm for full nuclei and 2 μm for magnified panels.

Ito et al., Figure S8

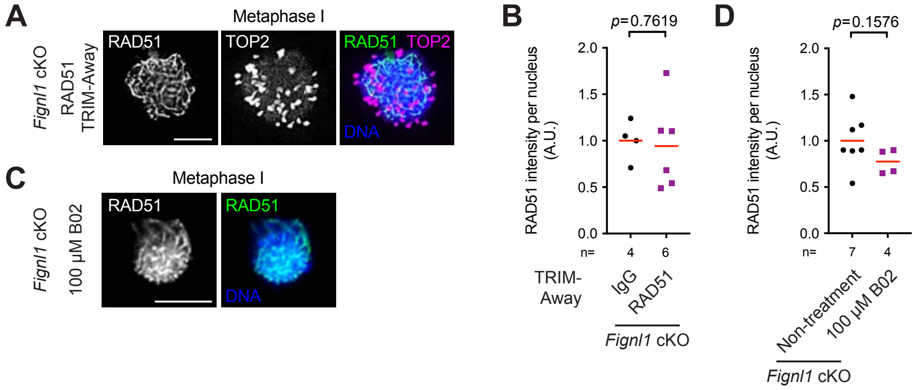


**Fig. S8. Chromosomal RAD51 binding resistant to RAD51 TRIM-Away and chemical inhibitor in metaphase-I *Fignl1*-deficient oocytes.**

(A) Representative images of chromosome spreads of RAD51 TRIM-Away oocytes at metaphase I stained with DAPI (DNA, blue) and immunostained for RAD51 (white in the left panel and green in the right panel) and TOP2 (white in the middle panel and magenta in the right panel) from *Fignl1* cKO mice.

(B) Signal intensities of chromosomal RAD51 on chromosome spreads of control (black circles) and RAD51 (purple squares) TRIM-Away oocytes at metaphase I from *Fignl1* cKO mice. The red bars are means.

(C) Representative images of fixed metaphase-I oocytes treated with B02 RAD51 inhibitor stained with DAPI (DNA, blue) and immunostained for RAD51 (white in the left panel and green in the right panel) from *Fignl1* cKO mice. GV oocytes were cultured with 100 μM B02 for 7 h and fixed. Images are optical slices to show the middle of a nucleus.

(D) Signal intensities of nuclear RAD51 in fixed metaphase-I oocytes cultured without (black circles) and with (purple squares) 100 μM B02 from *Fignl1* cKO mice. The red bars are means.

The results of two-tailed Mann-Whitney *U*-tests are indicated in the graphs. Total numbers of oocytes analyzed are indicated below the graphs. Genotypes of indicated animals are: *Fignl1* cKO, *Fignl1*$^{flox/\Delta}$ *Stra8-Cre*$^{+}$. Scale bars in (A) and (C), 10 μm.

Ito et al., Figure S9

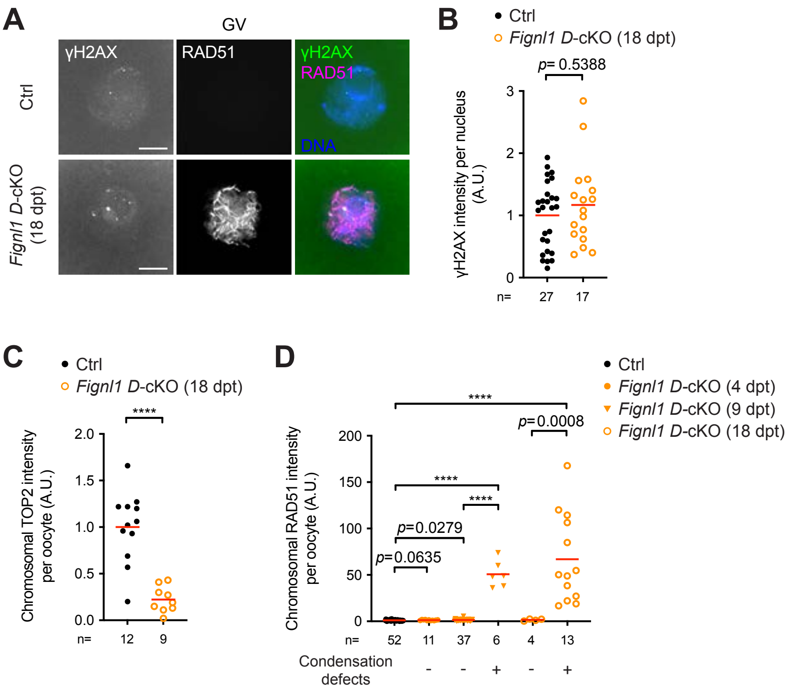


**Fig. S9. Aberrant RAD51 accumulation and chromosome condensation defects with little increase in DNA damage by *Fignl1* depletion in growing oocytes.**

(A) Representative images of fixed GV oocytes stained with Hoechst 33342 (DNA, blue) and immunostained for γH2AX (white in the left panels and green in the right panels) and RAD51 (white in the middle panels and magenta in the right panels) from Ctrl and *Fignl1 D*-cKO mice at 18 days post-tamoxifen treatment (dpt). Images are optical slices to show the middle of nuclei. Scale bars, 10 μm.

(B) Signal intensities of nuclear γH2AX in fixed GV oocytes from Ctrl (black circles) and *Fignl1 D*-cKO (orange open circles) mice at 18 dpt. The red bars are means.

(C) Signal intensities of chromosomal TOP2 on chromosome spreads of metaphase-I oocytes from Ctrl (black circles) and *Fignl1 D*-cKO (orange open circles) mice at 18 dpt. The red bars are means.

(D) Signal intensities of chromosomal RAD51 on chromosome spreads of metaphase-I oocytes from Ctrl (black circles) and *Fignl1 D*-cKO (orange filled circles, triangles, and open circles at 4, 9, and 18 dpt, respectively) mice. Oocytes with chromosome entanglement or mass are classified as positive for condensation defects. The red bars are means.

The results of two-tailed Mann-Whitney *U*-tests are indicated in the graph: ****$p$

≤ 0.0001. Total numbers of oocytes analyzed are indicated below the graph. Genotypes of indicated animals are: Ctrl, *Fignl1$^{flox/Δ}$*; *Fignl1 D*-cKO, *Fignl1$^{flox/Δ}$ Dppa3-MCM$^{+}$*. In (B) and (C), oocytes with linear RAD51 immunostaining were selectively analyzed for *Fignl1 D*-cKO mice.

**Movie S1.** Related to Fig. 2A. Time-lapse live-cell imaging of histone H2B–mRFP and EGFP–MAP4MTBD in a Ctrl GV oocyte following dbcAMP release for meiotic resumption and maturation. Fluorescence images are shown as maximum intensity *z*-projections, accompanied by single-plane bright-field images. Images were acquired at 5-min intervals until 2 h 10 min and at 20-min intervals thereafter.

**Movie S2.** Related to Fig. 2A. Time-lapse live-cell imaging of histone H2B–mRFP and EGFP–MAP4MTBD in a *Fignl1* cKO GV oocyte following dbcAMP release for meiotic resumption and maturation as in Movie S1.
